# Reptilian hunchbacks: a systematic review and meta-analysis of spinal deformities in wild reptiles

**DOI:** 10.1101/2025.06.23.661054

**Authors:** Gergely Horváth

## Abstract

Deformities of the spinal column—such as kyphosis, scoliosis, lordosis, and their various combinations—are known to affect all major vertebrate taxa. In reptiles, these deformities are relatively common in captivity, but reports from wild populations are also frequent. Since the 2000s, the number of published cases and affected species has increased considerably. However, most observations still appear as natural history notes or brief reports, often treating spinal deformities as mere curiosities without examining their aetiology, pathology, or prevalence. As a result, we know virtually nothing about the potential ecological impacts of spinal deformities on the life history of affected individuals or populations. Here, based on a systematic review of peer-reviewed and grey literature, complemented by unpublished observations gathered through standardized social media inquiries, I present the most comprehensive database and review of spinal deformities in wild reptiles to date. I identified 351 observations from 139 sources, covering 103 species across 24 families and 35 countries. Unpublished observations accounted for 13.3% of the database. Spinal deformities appear across all major reptile clades, though Testudines are clearly overrepresented (59.8%), likely due to their greater popularity, visibility, and research attention compared to other groups. Prevalence data were available or could be extracted from 51 records (36 species). A phylogenetic meta-analysis revealed a global prevalence effect size of 0.23%. However, no significant effects of phylogeny, habitat use strategy, or habitat type were detected—though this may reflect limitations of the dataset. Kyphosis appeared more frequently in (semi)aquatic Testudines than in primarily terrestrial squamates, whereas the opposite pattern was observed for scoliosis. Nonetheless, the underrepresentation of (semi)aquatic squamate taxa in the dataset urges caution when interpreting these phylogenetic inferences. Spinal deformities may substantially impact key life-history traits such as survival, growth, and reproduction, but direct evidence remains scarce and mostly anecdotal. To enhance ecological relevance, future studies should adopt more detailed and standardized reporting— incorporating occurrence frequency, habitat characteristics, morphometrics, imaging data, and assessments of behavioural or locomotor limitations.

## I. INTRODUCTION

Spinal column deformities are observed in every major vertebrate group, with reported cases in fish (e.g., Ayed et al., 2007; Jawad & Ibrahim, 2018), amphibians (e.g., Martínez et al., 1992; Gamble et al., 2005), birds (e.g., Kentaro et al., 2005) and mammals (e.g., Berghan & Visser, 2000; Hata et al., 2020). Unsurprisingly, reptiles are no exception. The oldest known record of spinal deformities comes from a Permian aquatic parareptile, *Stereosternum tumidum*, exhibiting scoliosis (lateral deviation of the vertebral column) (Szczygielski et al., 2017). Other common forms of these deformities include kyphosis (abnormal increase of the posterior convexity of the vertebral column), and lordosis (abnormal concave curvature of the vertebral column). Deformities can also occur in combination, with the most common being kyphoscoliosis (Fig. 1), while the simultaneous presence of all three conditions is referred to as rhoecosis (cf. Smith & Fitzgerald, 1983; de Carvalho et al., 2021).

**Figure 1.**
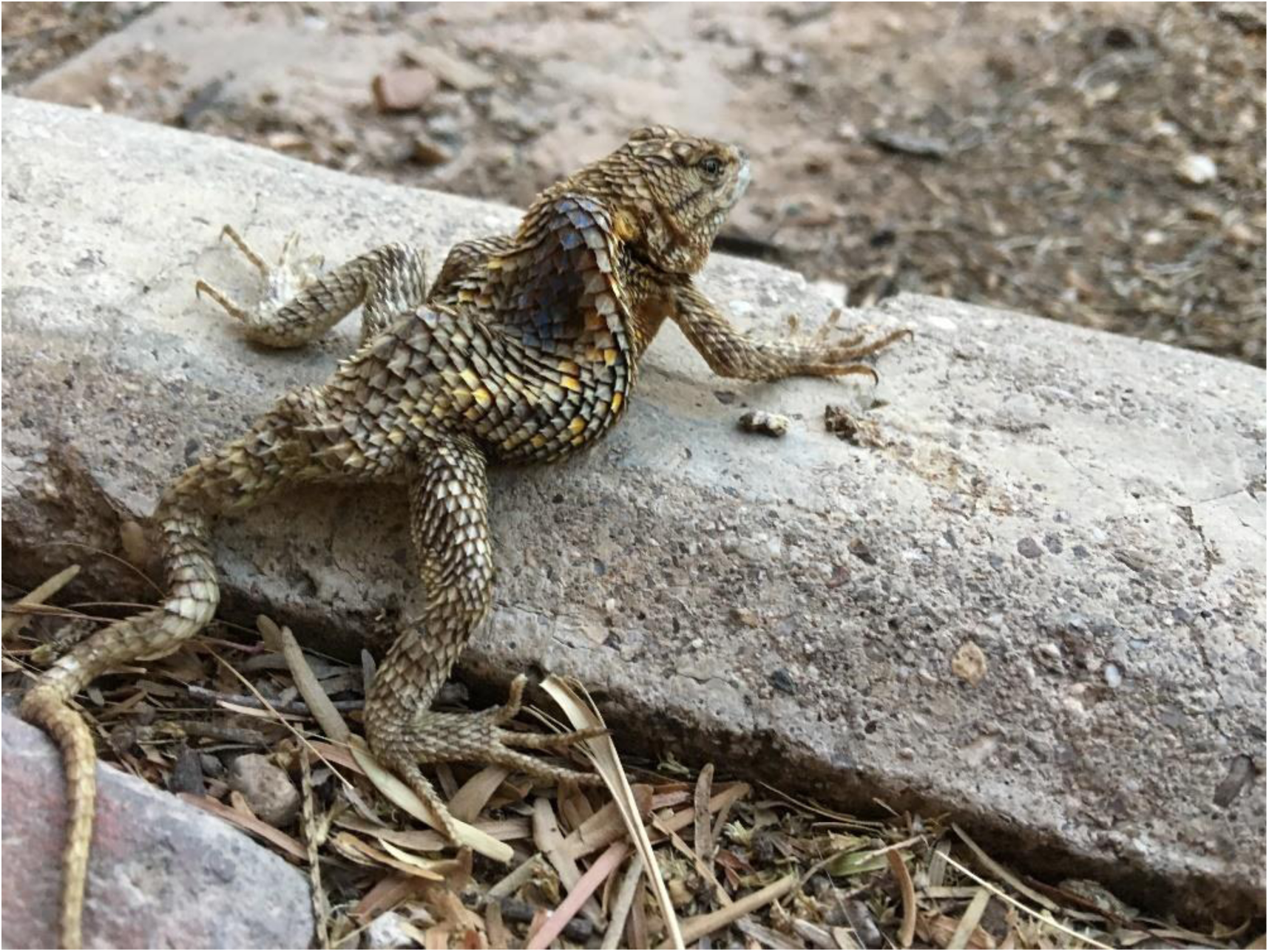
Kyphoscoliotic desert spiny lizard (*Sceloporus magister*). Photograph courtesy of Dennis Caldwell.

The earliest scientific reports of spinal deformities in reptiles date back to the early 1900s, with all initial cases involving Testudines. Wandolleck (1904) offers a detailed examination of a kyphotic Hermann’s tortoise (*Testudo hermanni*) from the Dresden Zoo, though he refrains from speculating on the exact cause of the deformity. Similarly, Vogt (1922), Gressit (1936a, b, c), Mertens (1940), and Necker (1940) describe their observations of ‘humpbacked’ specimens as curiosities. Although some of these authors mention factors such as environmental contamination and growth disorders as potential causes of the deformities, they fail to provide a broader context for the pathology or discuss the potential impact of the condition on the affected individuals. While the aetiology and pathogenesis of these conditions still remain unclear, spinal deformities appear to be highly multifactorial, with no single aetiopathological cause that can be fully accounted for. Importantly, our understanding of the causes and development of spinal deformities is largely based on human medicine, but due to the anatomical similarities of the spine across vertebrates, animal models are frequently studied (Janssen et al., 2011; Oullet & Odent, 2013; Lv et al., 2021). However, despite structural similarities, certain forms of spinal deformities seem to be taxon specific. For example, lumbar lordosis and idiopathic scoliosis are conditions predominantly associated with humans, likely as a consequence of bipedal posture (cf. Kouwenhoven & Castelein, 2008; Sparrey et al., 2014). Additionally, the factors contributing to spinal deformities, as well as the mechanisms of their development, may vary across different taxa.

### (1) The most common factors behind spinal deformities in reptiles

Although physical trauma and vertebral fractures were previously suggested as primary causes of spinal deformities, and such injuries can indeed lead to kyphosis or scoliosis, occurrences are rather sporadic or secondary (see Smith & Fitzgerald, 1983; Feltrin et al., 2009). In contrast, pathological fractures resulting from nutritional or metabolic diseases are more likely to cause spinal deformities (see below). Perhaps the most diverse group of aetiological agents for kyphosis and scoliosis has been proposed in case of Testudines. Early authors suggested that these deformities emerge due to differential growth rates of the skeletal components of the carapace, possibly in combination with the premature fusion of the vertebral column with the carapacial plates (see Smith 1947). Based on observations of hatchlings, Cagle (1950) and Williams (1957) proposed that abnormal yolk retraction could lead to early skeletal fusion of the carapace and, consequently, a malformed spine. However, this idea was not further tested, although it has been frequently referenced. Another possibility is that spinal deformities have a neoplastic origin (cf. Johnson, 2012; Mitchell & Tully, 2016), though cases that clearly attributable for abnormal tumour growth have not been documented. As in other vertebrates, spinal deformities in reptiles are typically multifactorial abnormalities and often represent just one clinical sign within a complex disease profile. Given the relative prevalence of spinal deformities in captive reptiles, advancements in reptile veterinary medicine have allowed for the identification of key contributing factors (cf. Divers & Stahl, 2019).

#### (a) Anomalies of embryonic development

In reptiles, kyphosis, scoliosis, lordosis, and their combinations are commonly reported congenital malformations. A study by Sant’Anna et al. (2013) found that spinal deformities were the most frequent malformations in newborn Jararaca (*Bothrops jararaca*) and Cascadel rattlesnake (*Crotalus durissus*). Although the overall prevalence of all malformations was negligible (2.3%), among malformed neonates, kyphosis alone was accounted for more than 60% of the cases in both species (*B. jararaca*: 67.4%, *C. durissus*: 75%). In contrast, kyphosis was less common in sea turtles; for example, in olive ridley sea turtle (*Lepidochelys olivacea*) kyphosis was present in only 13.2% of malformed embryos or neonates (Bárcenas-Ibarra & Gasca, 2009), with some studies reporting even lower prevalence rates (cf. Bárcenas-Ibarra et al., 2015, 2017). Importantly, although spinal deformities often appear as standalone conditions, in many cases they are combined with other deformations, e.g., cranial malformations, anophthalmia, buphthalmia, coiled tail, schistogastria (see Bell, Spotila & Congdon, 2006; Sant’Anna et al., 2013; Carvalho, 2014; Elsey & Stelly, 2018). These multifactorial defects are not necessarily fatal, and if they do not interfere with locomotion and feeding, individuals might reach adulthood (see Schachner et al., 2017).

Identifying the primary causes of congenital spinal deformities is challenging due to the intricacy of vertebrate embryonic development. This highly synchronized biological process is vulnerable to aberrations and errors at various stages including gametogenesis, fertilization, blastogenesis, embryogenesis, and fetogenesis (cf. de Carvalho et al., 2021). While many congenital spinal deformities have a genetic origin, suboptimal environmental factors also play a significant role. Unfortunately, the interaction between genetic and environmental factors remains poorly understood, making it difficult to attribute developmental anomalies to a single cause. Regarding genetic underpinnings, geographic isolation and population fragmentation are perhaps the most significant factors contributing to the restriction of the genetic pool in natural populations. Inbreeding and genetic drift along with the subsequent expression of genes associated with developmental anomalies, often results from this restriction. Thus, in small and isolated populations the frequency of skeletal deformities is found to be increased compared to those with high genetic variability (cf. Madsen, Stille & Shine, 1996; Daltry et al., 2001; Lindsay et al., 2020).

The influence of agents originating from both the external and maternal environments can significantly impact embryonic development in both oviparous and viviparous reptiles, as they both employ a simple form of placentation (Blackburn & Flemming, 2009). Embryos of several oviparous reptile taxa are vulnerable to extremes in moisture, gas exchange and incubation temperature. Insufficient environmental humidity is frequently cited and investigated as key teratogen in abnormal shell development in Testudines, often leading to spinal deformities. Lynn and Ullrich (1950) found that suboptimal soil moisture during egg incubation can produce various abnormalities of the carapace of freshwater turtles, such as the painted turtle (*Chrysemys picta*) and the common snapping turtle (*Chelydra serpentina*).

Regarding extreme temperatures, Idrisova (2018) found that both grass snakes (*Natrix natrix*) and sand lizards (*Lacerta agilis*) incubated at high temperatures (29 °C and 34 C °) more frequently exhibited a wide range of deviations, including spinal deformations. Additionally, anomalies were more pronounced in these hatchlings, resulting in reduced survival. Similarly, Telemeco et al. (2013) reported that prolonged exposure to extreme incubation temperature increases the frequency of shell and scute abnormalities in painted turtles (*C. picta*). This is significant, as in Testudines, abnormal shell development can often lead to kyphosis (cf. Johnson, 2012; Juan-Sallés & Boyer, 2021)

Perhaps the most important emerging group of environmental teratogens for reptiles are pollutants (cf. Dolk & Vrijheid, 2003; Orós et al., 2021; Garcés & Pires, 2023). Since the 1990s, research on environmental toxicology in reptiles has increased, though it still receives less attention compared to contaminants in other vertebrates. Nonetheless, it is now clear that anthropogenic activities are seriously affecting natural reptile populations globally through various chemical substances, including heavy metals, pesticides, and certain drugs.

Considerable data suggest that in contaminated and highly urbanized areas, the incidence of congenital abnormalities is significantly higher (Gray et al., 2001; Gray, Smith & Chiszar, 2003; Bell et al., 2006; Martín-del-Campo et al., 2021).

#### (b) Nutritional and metabolic diseases

In captivity, the most common cause of skeletal spinal abnormalities is metabolic bone diseases (MBDs). MBD is not a single condition but rather an umbrella term that encompasses a group of related conditions that affect the integrity and function of bones (cf. Hedley, 2012a; Knafo, 2019; Juan-Sallés & Boyer, 2021). The most common MBD is nutritional secondary hyperparathyroidism (NSHP), although renal secondary hyperparathyroidism (RSHP) can also occur. NSHP typically arises from inadequate UV-B radiation, an improper temperature range, or insufficient dietary calcium or vitamin D3 (Hedley, 2012a; Carmel & Johnson, 2018; Knafo, 2019). While NSHP can potentially affect all reptile taxa, it is more prevalent in heliothermic lizards and Testudines. These reptiles, due to their mostly insectivorous and herbivorous diets, cannot rely solely on dietary sources of vitamin D3; thus, the activation of dietary provitamin D precursors via UV-B radiation is essential. Typical clinical signs of MBDs include pathological fractures, curvature of the spine, or deformed carapace in Testudines, and many bones appearing more curved than normal (Hedley et al., 2012b; Knafo, 2019; Juan-Sallés & Boyer, 2021).

It is noteworthy to mention a condition affecting Testudines known as shell pyramiding (Juan- Sallés & Boyer, 2021). Pyramiding refers to the abnormal growth of the scutes and the underlying bones of the carapace and is considered a distinct form of nutritional and metabolic disease. In affected specimens, material deposition between the scutes accelerates scute growth at a rate faster than the expansion of the underlying bone. This forces the keratin and bone to grow conically upward, causing a separation between the vertebrae and the carapace, with only extremely thin bone bridging from the dorsal vertebral body to the carapace. This condition commonly results in kyphosis, scoliosis, and lordosis. Pyramiding is thought to be a multifactorial process, likely caused by an imbalanced calcium-to-phosphorus ratio, increased growth rates, and low humidity (Weisner & Iben, 2003; Heinrich & Heinrich, 2016).

In contrast to lizards and Testudines, snakes appear to be less affected by MBDs because they consume their prey whole, providing sufficient dietary vitamin D3 and calcium. Additionally, while MBD and shell pyramiding are very common in captivity, these conditions are believed not to affect wild populations (cf. Carmel & Johnson, 2018; Juan-Sallés & Boyer, 2021).

However, their potential presence in natural populations cannot be entirely ruled out.

#### (c) Infection-induced osteophaty

Bacterial osteomyelitis and osteoarthritis, both being severe bone infection, are common in reptiles and can lead to stiffness, pathological fractures, and spinal deformities when the spinal column is affected (Rasche et al., 2022). Vertebral osteomyelitis and osteoarthritis are especially common in snakes (e.g., Isaza, Garner & Jacobson, 2000; Ramsay et al., 2002; Schröter et al., 2005). The bacteria most commonly associated with this condition are Gram-negative rods including *Citrobacter* sp., *Salmonella* sp., *Proteus* sp., *Pseudomonas* sp., and *Escherichia coli* (cf. Ramsay et al., 2002; Kwon et al., 2020). Gram positive bacteria seems to be rarely caused osteomyelitis in reptiles, however, there are documented cases involving *Staphylococcus* sp., *Streptococcus* sp., and *Enterococcus* sp. (cf. Isaza et al., 2000; Schröter et al., 2005). On rare occasions, fungi have been reported as causative agents of bone infection (cf. Heatley et al., 2001).

Bacteria often invade the spinal column through direct extension, typically resulting from wound contamination or trauma to nearby soft tissues. In the case of *Salmonella* sp., a common inhabitant of the reptile digestive tract, bacterial spread can occur during periods of immune suppression. This may lead to septicaemia, allowing the bacteria to reach the spinal column via hematogenous dissemination (cf. de Souza et al., 2014). Some researchers have proposed that osteomyelitic spinal lesions could be caused by bacterial endotoxins (see in Isaza et al., 2000). For a definitive histological diagnosis of the bacterial origin of osteomyelitis and osteoarthritis, isolating bacteria from bone cultures is essential. This requirement may explain why reports in the literature almost exclusively involve captive animals. However, Isaza et al. (2000) observed a strong correlation between bone culture and blood culture results, suggesting that blood cultures, being a simpler and far less invasive method, could be a feasible alternative for diagnosing active bacterial osteomyelitis and osteoarthritis.

### (2) Ecological implications

It is important to consider the impact of potentially damaging malformations, such as spinal deformities, on the lives of individual reptiles and, ultimately, their populations. Despite an increasing trend in observations, spinal deformities remain relatively rare in wild populations. It has been suggested that this rarity may result from the higher mortality rate of affected individuals (cf. Garín-Barrio et al., 2011). However, numerous observations report adult individuals with spinal deformities that appear to experience no significant negative effects on foraging or mobility (e.g., Garín-Barrio et al., 2011; Avila et al., 2013; Pérez-Delgadillo et al., 2015; Domínguez-De la Riva & Carbajal-Márquez, 2016; Valdez-Villavicencio, Hollingsworth & Galina-Tessaro, 2016; Horváth, Martín & Herczeg, 2021). This suggests that spinal deformities may not necessarily reduce the survival chances of affected individuals.

Furthermore, Mitchell and Johnston (2014) demonstrated that the growth pattern of a Florida chicken turtle (*Deirochelys reticularia chrysea*) with kyphoscoliosis was comparable to that of individuals without abnormalities within the same population.

Another important yet unresolved question is whether habitat deterioration and pollution contribute to an increased frequency of spinal deformities. Although no conclusive link has been established between the development of spinal deformities and contaminants, the global presence of pesticides, herbicides, fungicides, and petrogenic or pyrogenic polycyclic aromatic hydrocarbons (PAHs) poses potential risks even to (quasi-)natural populations (cf. Alegbeleye, Opeolu & Jackson, 2017). Supporting this concern, there are observations of spinal malformations in which agrochemicals have been suggested as possible causative agents (Pérez-Delgadillo et al., 2015; Ramírez-Jaramillo, 2018).

Finally, it remains largely unknown whether different habitat use strategies (i.e., terrestrial, arboreal, aquatic, and semi-aquatic) make reptiles more prone to spinal deformities in general, or if certain types of deformities are more frequently associated with specific strategies. For instance, it is suggested—though not conclusively demonstrated—that kyphosis primarily affects species adapted to aquatic environments (e.g., turtles and crocodiles; cf. Boede & Sogbe, 2000; Tucker, Lamer & Dolan, 2007), while scoliosis appears more common in terrestrial species (e.g., snakes and lizards; cf. Grogan, 1976; Simbotwe, 1983; Frutos et al., 2006).

### (3) Aims of the review

Reports of spinal deformities in wild reptiles have become common over the past two decades. The number of published cases, as well as the range of affected species, has noticeably grown since the 2000s. However, it remains unclear whether this rise in reports reflects a genuine increase in deformities or is simply the result of heightened research interest in these “hunchbacks.” Alternatively, the intensification of key environmental stressors, such as human-induced environmental changes and pollution, could be driving this trend. Unfortunately, most observations treating spinal deformities as mere curiosities without investigating their aetiology, pathology, or prevalence. Consequently, we know virtually nothing about the potential ecological impacts of spinal deformities on the life history of affected individuals or populations.

The primary objective of this study is to present the first comprehensive review of spinal deformities in wild reptiles, based on both published reports and previously unpublished personal observations. Emphasis was placed on assessing the occurrence of various types of spinal deformities across reptile families. Where possible, the prevalence of spinal deformities within populations was extracted or calculated. Prevalence data were analysed using a phylogenetic meta-analysis to estimate a global effect size for the frequency of spinal deformities and to evaluate whether factors such as habitat type (natural, seminatural or urbanized) and habitat use strategies (aquatic, semiaquatic, terrestrial, arboreal) influence prevalence. These findings are used to discuss the potential ecological impacts of spinal malformations and to provide recommendations for future research on this phenomenon.

## II. METHODOLOGY

I compiled records on spinal deformities in reptiles from published, peer-reviewed literature, as well as non-peer reviewed (‘grey’) literature, spanning the earliest available material (1904) to January 2025. Given that most records consist of single-observation natural history notes, short communications, or research articles, I systematically reviewed the relevant sections of the complete published issues of the following journals: *Boletín de la Associacón Herpetológica Española* (1990-2024), *Bulletin of the Chicago Herpetological Society* (1990- 2023), *Herpetology Notes* (2014–2025), *Herpetological Review* (1967–2024), *Herpetozoa* (2019-2024), *The Herpetological Bulletin* (1980–2024), *The Herpetological Journal* (1985–2025), *Mesoamerican Herpetology* (2014–2017), *Salamandra* (1965-2024), *Reptiles & Amphibians* (2009-2025); *Revista Española de Herpetología* (1996-2010); *Revista Lationoamericana de Herpetología* (2018-2024) and *Russian Journal of Herpetology* (1994- 2024).

Additionally, online libraries and databases, i.e., *Google Scholar* and *Web of Science* were queried using relevant search terms: ‘kypho’, ‘scolio’, ‘lordo’, ‘spine’, ‘spinal’, ‘deform’, and ‘malform’. These terms were also searched in German, Spanish, Portuguese and French. To identify photographic records, I searched platforms such as *Flickr* and *Google Images*. Any records cited in short notes as containing pertinent information were retrieved and reviewed. For materials unavailable online, including hard-copy journals or books, requests were made to libraries or individual researchers. Lastly, unpublished observations were solicited by circulating requests in several online herpetological groups (*California Herping*, *European Herping*, *Field Herping*, *Field Herpetology*, *Field Herping Europe*, *Field Herping North America*, *Herping Arizona*, *Herping Lao*, *Herping Southeast Asia*, *Herping in Thailand*, *Kentucky Herping*) on Facebook to gather novel records.

### (1) Database construction

I constructed a database by extracting the following information: order, superfamily, family, genus, scientific and common names, habitat use strategy (terrestrial, arboreal, aquatic, and semiaquatic), location of observation (country and specific locality), geographic region, and biogeographic realm (when specified). Additionally, SW4 coordinates and elevation were included when available, as well as the exact date of observation and the type of malformation (kyphosis, scoliosis, lordosis, kyphoscoliosis, kypholordosis, lordoscoliosis, and rhoecosis).

On some sporadic occasions, particularly in early 20th-century observations, no specific type of malformation was provided (e.g., Hildebrand, 1930; Gressit, 1936c), or incorrect terminology was used (e.g., Vogt, 1922; Ahboucha & Gamrani, 2001; Feltrin et al., 2009). In such cases, I reviewed the diagnosis based on photographic evidence. Diagnostic method (observation, radiography, and computerized tomography) was also recorded. When possible, morphometric data, including size (in mm; snout-to-vent length [SVL] or carapace length) and body weight (g), as well as sex, were extracted.

The prevalence of spinal deformities was calculated for records that reported observed cases, and the total number of individuals examined from the same population. In cases where multiple individuals with the same type of malformation from the same population were reported, I treated each deformed individual as a separate observation but used their total number to calculate prevalence. Therefore, the prevalence value is the same for these observations. Different types of malformations from the same populations were treated separately when calculating prevalence. If a record reported more than a certain number of individuals examined (e.g., >100 individuals), I used 100 as the total number for calculating prevalence.

The names of all reported species were cross-checked against the reptile database (Uetz et al., 2025) to obtain the most recent classification for each species, genus, and family, as well as the taxonomy and number of species within each family. All observations were classified to the species level.

### (2) Meta-analysis

Meta-analyses on proportions, such as prevalence, are commonly conducted, particularly in medicine (e.g., Pringsheim et al., 2014), epidemiology (e.g., Faustino et al., 2022), and clinical psychology (e.g., Tian-Ci Quek et al., 2019). In such meta-analyses, each study contributes a specific number of observed cases or successes along with a corresponding total sample size. In this analysis, I extracted 51 effect sizes, each with an associated total sample size, from 40 studies covering 36 species across 15 families (Supplementary Table S1).

Proportional data are often not centred around 0.5 and frequently exhibit significant skewness, deviating from a normal distribution (Wang, 2023). Using raw proportions as the effect-size metric in such cases can result in an underestimation of confidence interval coverage around the weighted average proportion and an overestimation of heterogeneity among observed proportions (Lipsey & Wilson, 2001). Consequently, assuming normality may introduce bias, potentially leading to misleading or invalid inferences (Feng et al., 2014; Ma, Chu & Mazumdar, 2016). To address this skewness, I applied a logit (log odds) transformation, converting the proportions into their natural logarithm using the following equation:

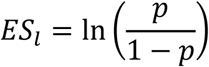

where *p* represents the proportion.

The meta-analytic approach assumes that variance between observations due to sampling error can be approximated by the squared standard error. However, sampling variance is higher for low-precision estimates, such as those based on a small sample size or a low number of repeats. This can result in (i) low-precision studies receiving disproportionate weight, (ii) an overestimation of biological variation, and (iii) potential bias in the mean effect size if publication bias is more prevalent among low-precision studies. Sampling variance was calculated using the following equation (following Wang, 2023):

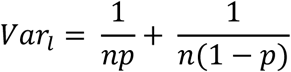

where *p* represents the proportion and *n* is the total sample size.

Preliminary descriptive analyses revealed a strong association between study identity and species. Except for Rhodin et al. (1984), Barcanes-Ibarra et al. (2015), Mitchell et al., (2019), and Jackson & Zappalorti (2020) all studies focused on a single species, while some species were observed by multiple studies (*Caretta caretta*, N = 2; *Dermochelys coracia*, N = 2; *Lepidochelys olivacea*, N = 4; *Crysemys picta*, N = 5; see Supplementary Table S1). I added species ID to account for sources of non-independence arising from effect sizes coming from the same species. In the same time, effect size ID was added to account for variation in effects across individual effect sizes.

Biological and environmental differences between studies are expected to affect the variation of association between components of behavioural strategy, thus, I accounted for these potential sources of heterogeneity by extracting and examining the effect of two moderator variables. The following variables were extracted from each study: (i) habitat use strategy (aquatic, semiaquatic, terrestrial, arboreal) and (ii) habitat type (natural, seminatural or urban [in the case of seven study, habitat type was not applicable]). Habitat type was determined through visual inspection of the surroundings of the SW4 coordinates using Google Maps.

Urban habitats were defined as city parks, ponds, or riverbanks located within city limits, whereas seminatural habitats included similar features situated in less frequented rural areas or outside city limits but still potentially subject to substantial human influence.

Information on the phylogenetic history of species was obtained based on published phylogenies available through the ‘Open Tree of Life’ (Hinchliff et al., 2015) via the *rotl* package (Michonneau, Brown & Winter, 2016). Taxon names were matched to records in the Open Tree Taxonomy to obtain relationships between species. It should be noted that, for compatibility with the OTL database, the Suwannee snapping turtle (*Macrochelys suwanniensis*) was treated as a synonym of the alligator snapping turtle (*M. temminckii*).

However, this does not significantly affect the interpretation of the results. Due to the diversity of species in this meta-analysis, accurate estimation of branch lengths was not possible, thus, branch lengths were computed based on topology (see Supplementary Fig. S1) using the *ape* package (Paradis, Claude & Strimmer, 2004) in R 4.5.1. (R Developmental Core Team, 2025). Phylogenetic heritability or phylogenetic signal (H^2^) was calculated as the proportion of total variance in effect size that can be explained by phylogenetic variance (Hadfield & Nakagawa, 2010), equivalent to Pagel’s λ (Pagel, 1999). H^2^ = 0 indicates no phylogenetic relatedness among effect sizes (Lynch, 1991).

I conducted the meta-analysis using a random-effects model, estimated by the restricted maximum likelihood method in the R package *metafor* (Viechtbauer, 2010). I first ran intercept-only mixed model (with random effects) to determine the mean effect size across all studies. To estimate heterogeneity of effect sizes, I used I^2^ statistics (e.g., Horváth, Garamszegi & Herczeg, 2023; Mizuno et al., 2024; Lin, 2020) modified for multilevel meta-analytic models (Nakagawa & Santos, 2012): total heterogeneity (I^2^_total_) was partitioned into phylogenetic variance (I^2^phylogeny), species ID variance (I^2^_species_), study ID variance (I^2^_study_) and residual variance (I^2^_residual_). Low, moderate and high heterogeneities refer to I^2^ of 25%, 50% and 75%, respectively (Higgins & Thompson, 2002). Next, I constructed a series of meta-regression models to identify the most important moderators (see above) (Horváth et al., 2023). As the sample size was somewhat limited, I chose to avoid complex models, instead, I conducted fixed-effect mixed models to estimate the mean effect size for each moderator separately (Horváth et al., 2023; Winternitz et al., 2017).

## III. RESULTS

### (1) Overview of published cases

The search yielded 139 eligible sources, from which 351 observations of spinal deformations were extracted across 24 families, 63 genera, and 103 species (see Supplementary Table S1). The oldest observation dates back to 1898: a specimen of *Crocodylus acutus* found dead at an undisclosed location in Panama (Rossmann, 2001). The overwhelming majority of observations came from peer-reviewed literature (304 records, 86.6%), including journal articles (237 records, 67.5%). A single dissertation (Carvalho, 2014) contributed 61 records (17.4%), while books accounted for 6 records (1.7%). Online social and popular media platforms provided 10 records (2.8%), and grey literature contributed 37 records (10.5%), all of which were unpublished observations. The countries contributing the most records are the USA (154 records, 43.9%), followed by Brazil (65 records, 18.5%; note that without the dissertation of Carvalho [2014] the share of Brazil changes significantly: 4 records, 1.1%) and Mexico (46 records, 13.1%) (Fig. 2). While all observations could be linked to a country, the specific locality was provided or could be inferred for less than half of them (165 records, 47%). Exact coordinates were available in only 83 cases (23.6%), and elevation was recorded in 45 cases (12.8%).

**Figure 2.**
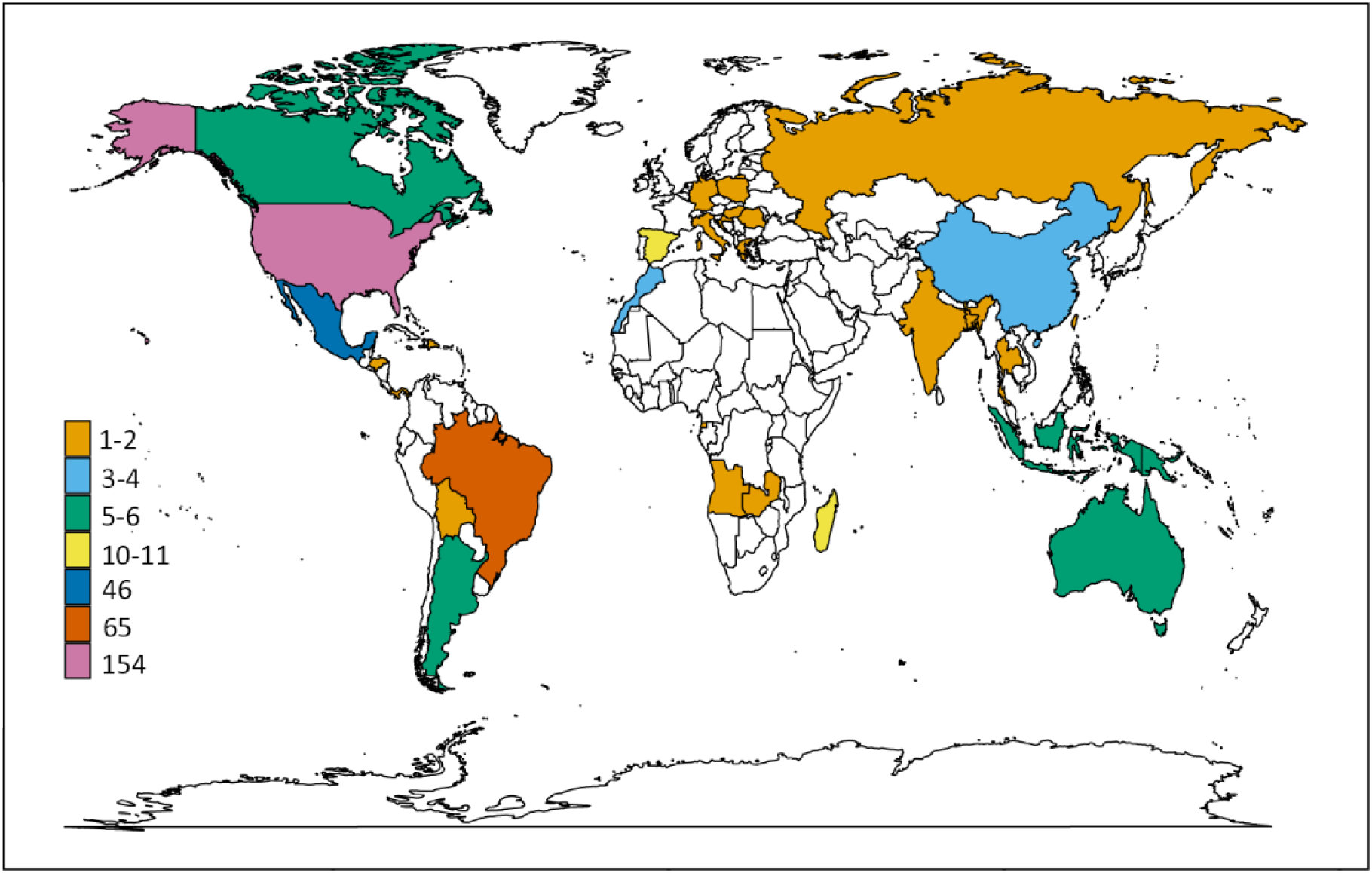
Global distribution of spinal deformation observations in wild reptiles.

More than half of the spinal deformations were cases of kyphosis (202 records, 57.5%; in five cases, the exact diagnosis was questionable) (Fig. 3). Scoliosis was reported in 64 cases (18.2%; ten cases were questionable), and in one instance it was not possible to distinguish between scoliosis and kyphosis. Kyphoscoliosis was indicated in 63 cases (17.9%; four cases were questionable). Only 3.7% of cases involved lordosis (13 cases, one of which was questionable). Rhoecosis was reported in five cases (1.4%), kypholordosis in two cases (0.6%), and there was a single case of lordoscoliosis (0.3%). In most cases, diagnoses were made by visual observation of the specimens. Imaging techniques—such as radiography and computer (micro)tomography—were used to diagnose spinal malformations in 92 cases (26.2% of all observations), reported in 27 studies (19.8% of all studies). Some form of additional information about the affected specimens was available in most cases. Sex was reported for 201 observations (57.2%), age for 232 observations (66.1%), size (snout–vent length or carapace length) in 164 cases (46.7%), and body mass in 72 cases (20.5%). The origin of vertebral deformations (congenital *vs.* postnatally acquired) was reported or could be extracted for 133 records (37.8%), while possible cause of the observed malformations was suggested by the author(s) in 94 cases (26.7%). Information on whether or not the malformations affected locomotion could be gathered in 67 cases (19%).

**Figure 3.**
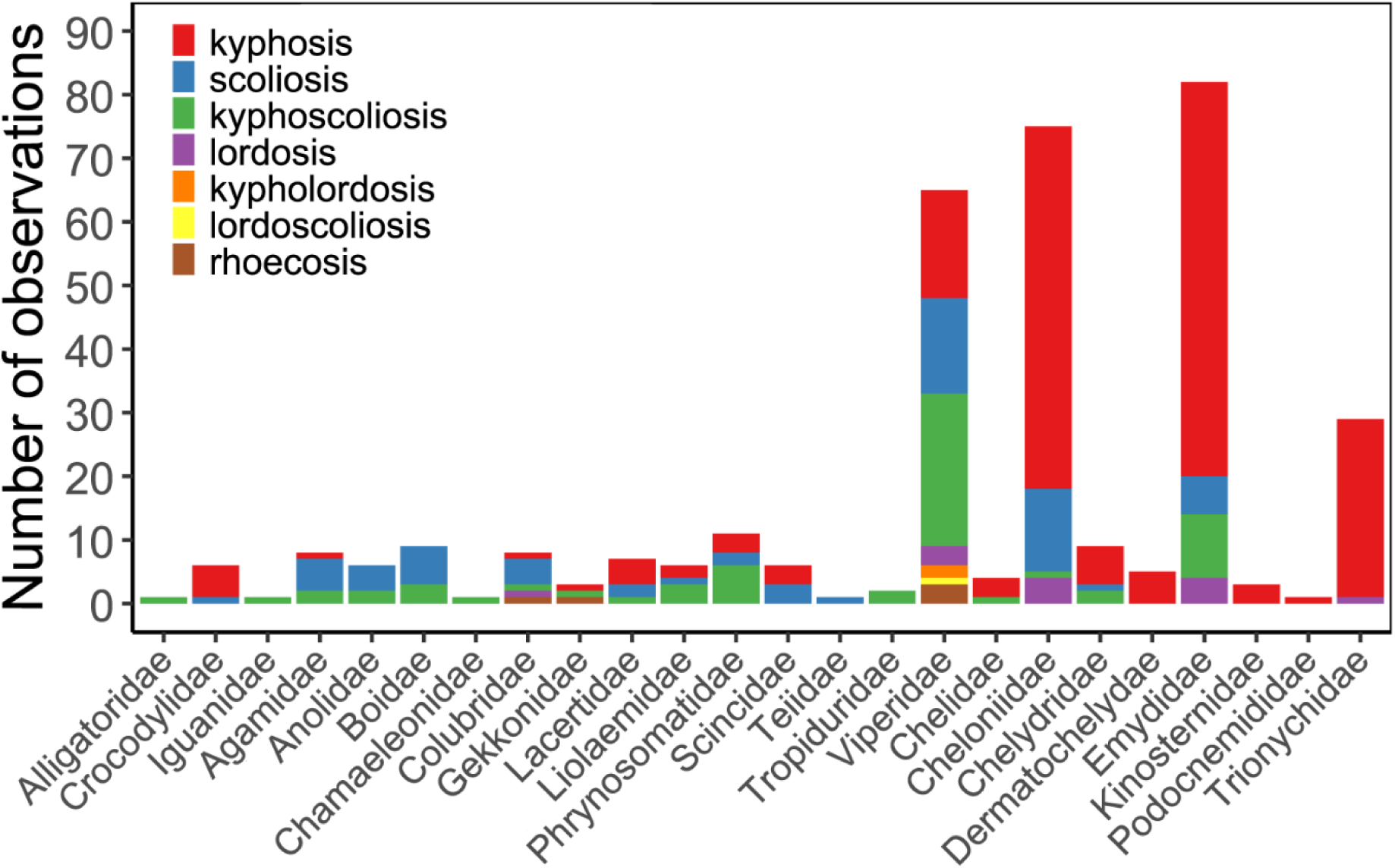
Types of vertebral malformation observations in different reptile families according to peer-reviewed and ‘grey’ literature publication records.

### (2) Prevalence of spinal malformations

The distribution of malformation types among the 51 effect sizes closely mirrored that of the full dataset: 32 cases (62.7%) were kyphosis, nine (17.6%) scoliosis, eight (15.7%) kyphoscoliosis, and two (3.9%) lordosis.

Based on the phylogenetically controlled meta-analysis, the overall mean effect size— calculated as the log odds of proportions—was statistically significant, indicating a mean malformation prevalence of 0.23% (this value is the back-transformed estimate from the log odds; see the provided R code for details) (estimate = -6.08, 95% CI = -6.57 to -5.59, t(df = 50) = -25.24, p < 0.001). Total heterogeneity across effect sizes was high (I²total = 80.12%) and was explained mostly by study differences (I²study = 71.57%). Effect size ID, representing residual variation was relatively low (I²residual = 8.54%), while both phylogenetic and among species differences were negligible (I^2^phylogeny & I^2^species <0.001%). In line with this, heritability was negligible too (mean H^2^ = 0.89).

On average, prevalence for aquatic taxa was 0.21% (estimate = -6.19, 95% CI = -7.02 to - 5.35, t(df = 47) = -14.99, *P* < 0.001), 0.29% in semiaquatic taxa (estimate = -5.86, 95% CI = -7.48 to -4.24, t(df = 47) = -7.28, *P* < 0.001), 0.3% in terrestrial taxa (estimate = -5.82, 95% CI = -7.3 to -4.35, t(df = 47) = -7.93, *P* < 0.001) and 0.23% in arboreal taxa (estimate = -6.09, 95% CI = -8.49 to -3.69, t(df = 47) = -5.12, *P* < 0.001). When comparing habitat use strategies, none of the differences was statistically significant (Fig. 6a). On average, prevalence in natural habitats was 0.13% (estimate = -6.64, 95% CI = -7.36 to -5.93, t(df = 40) = -18.72, *P <* 0.001), 0.21% in seminatural habitats (estimate = -6.16, 95% CI = -7.57 to -4.74, t(df = 40) = -8.79, *P <* 0.001) and 0.21% in urbanized habitats (estimate = -6.15, 95% CI = -8.22 to -4.08, t(df = 40) = - 6.01, *P <* 0.001). Again, there was no statistical difference between habitat types (Fig. 6b)

The funnel plot showed no visual sign of funnel asymmetry (Supplementary Fig. S2a) and there was no detectable trend suggesting that more recent publications consistently showed lower or higher effect size values, which would have indicated the presence of time-lag publication bias (estimate = -0.009; 95% CI = -0.03 to 0.01, t(df = 49) = -0.71, *P* = 0.89; Supplementary Fig. S2b).

## IV. STUDY SYNTHESYS

### (1) Affected reptile families, global distribution and reporting

The observations were distributed across 24 families, representing one fourth of the 94 recognized reptile families (as of May 2025; Uetz et al. 2025). The proportion of affected families (relative to the total number of families in each group) was highest among crocodilians, at 66.6% (two out of three families), followed by Testudines (57.1%, eight out of 14), lizards (29.7%, 11 out of 37), and snakes (9.3%, three out of 32). No observations were recorded for tuataras or amphisbaenians (see Supplementary Table S1). It is perhaps not far-fetched to suggest that the total of 351 records is almost certainly a significant underestimate of vertebral malformations in reptiles. The frequency of recorded cases appears to be closely related to how easily a specific group of reptiles can be observed. For example, the relatively low proportion of affected snake families likely results from the fact that snakes are generally more secretive, solitary predators, often with low or nocturnal activity, whereas lizards and Testudines are more active during the day and typically occur at higher population densities (e.g., Maura et al., 2011). Fossoriality—a widespread evolutionary strategy among both lizards and snakes—is another factor that may contribute to the lack of observations: No cases of vertebral deformities were recorded from entirely or highly fossorial lizard taxa such as *Anguidae* and *Amphisbaenia*, while the proportion of affected families was extremely low in *Scincidae* (0.22%). In addition to observational biases, the perceived absence of vertebral deformities in fossorial lizards and snakes may also result from the fact that subterranean environments can buffer against certain environmental factors that contribute to malformations, such as climatic fluctuations and pathogens (cf. Cyriac & Kodandaramaiah, 2018).

Another key conclusion that can be drawn from the database is that observations are more common in taxa that are well-studied, popular among non-professionals, and in which deformities are clearly visible. Testudines generally meet these criteria, but perhaps most importantly: kyphotic and kyphoscoliotic individuals have a characteristic ’humpbacked’ appearance that is highly conspicuous even from a distance. All of these factors likely contribute to the significant overrepresentation of Testudines in the dataset. They account for 210 records in total—59.8% of all data—most of which come from peer-reviewed literature (181 records, or 51.6%). Among the novel, unpublished records gathered from internet sources or personal communications, Testudines make up a slightly higher proportion: 61.7% (29 records). The popularity and well-studied status of turtles among reptiles may explain why this group is not only overrepresented, but also why most extant families are affected by vertebral malformations. The proportion of affected species is particularly high in *Dermochelyidae* (100%) and *Cheloniidae* (66.6%)—perhaps the most studied and monitored groups of turtles (cf. Hays et al., 2024)—, but also in *Chelydridae* (60%), and *Emydidae* (46.6%). Interestingly, only three records coming from species that are considered terrestrial (*Terrapene carolina*, *T. ornata*, *T. triunguis*).

The dataset spans 35 countries (see Fig. 3), with the majority of observations coming from the Americas (287 records, 81.8%). Unsurprisingly, the United States provided the largest number of records (154), mainly due to two factors. First, 140 of these observations are turtle records, representing 66.6% of all turtle records in the dataset. This aligns with the fact that global turtle species richness peaks in the southeastern United States (Roll et al., 2017).

Regarding squamates, the United States contributes 14 records, the second-highest count after Brazil. However, if the dataset excludes Carvalho’s (2014) dissertation—which includes embryological deformities observed in offspring of wild-caught female jararaca (*B. jararaca*) and Cascabel rattlesnake (*C. durissus*)—the United States emerges as the leading contributor of squamate records as well. This pattern does not correspond with global biodiversity hotspots for lizards and snakes but is likely attributable to the large number of professional herpetologists and herpetology enthusiasts in the United States.

Overall, the records are not evenly distributed across the globe and do not reflect global reptile species richness. Although 146 records (41.5%) originate from the Global South, several biogeographical regions considered hotspots of reptile diversity are significantly underrepresented: the Afrotropical region (14 records, 5.4%), Australasia (11 records, 3.1%), and the Indomalayan region (12 records, 3.4%). This is particularly surprising when compared to studies using similar methodologies (e.g., Barr et al., 2020) and suggests that the dataset is biased by differences in language, culture, and scientific practices—factors known to hinder the publication or accessibility of certain records (Møller & Jennions, 2001; Amano, González-Varo & Sutherland, 2016; Barr et al., 2020). It should be noted that, although I made every effort to include all available records published in English, German, Spanish, Portuguese, French, and Russian, it is still possible that some non-English publications were missed.

Another factor that may contribute to the uneven global distribution of spinal deformation records is that such observations were—and to some extent still are—regarded as mere curiosities. This likely explains the limited number of peer-reviewed publications, typically no more than one or two per decade until the mid-1900s (Fig. 5). Notably, there has been a significant increase in both the number of reported cases and the diversity of affected species since the early 2000s (Fig. 5). Despite this growth, the majority of reports are still published in the form of natural history notes. For example, *Herpetology Notes* alone accounts for 72 records (20.5% of total observations and 30.3% of journal articles) across 40 species. While most of these publications lack detailed discussion of ecological implications, a few studies have begun to address this dimension (e.g., Bell et al., 2006; Mitchell & Johnston, 2014; Bateman et al., 2022; Najbar et al., 2022).

In the following sections, drawing on insights from the literature and the phylogenetic meta- analysis, I will address the most important questions concerning the ecology of vertebral deformities. Specifically, I will examine whether phylogeny and habitat use strategy influence the prevalence of deformities, whether environmental stress is a key factor behind their occurrence, and how such malformations affect behaviour, locomotion, growth, and survival. Finally, I will identify the most critical gaps in our current knowledge and outline potential directions for future research.

### (2) Does phylogeny and habitat use strategy affect prevalence?

Based on the results of the phylogenetic meta-analysis, differences across species (I^2^species) and phylogeny (I^2^phylogeny) appear to be of virtually negligible significance. This is supported by the estimate of phylogenetic heritability (H^2^), which indicates that only a small proportion of variance can be attributed to phylogenetic relatedness. However, it should be noted that the phylogenetic tree used in the analysis was constructed from a limited number of species and, more importantly, was strictly topological, lacking branch length information. Additionally, the results may be affected by limited statistical power in some taxa; notably, 80.3% of the prevalence estimates—41 records—came from turtles (26 species), therefore, the level of phylogenetic inertia detected in this study should be considered preliminary.

Vertebral malformation prevalence did not differ significantly among (semi)aquatic, terrestrial, and arboreal habitat use strategies (Fig. 4a). However, in the dataset, habitat use strategy is highly correlated with phylogeny, as all but one of the (semi)aquatic taxa are turtles. Therefore, this finding complements the previous result on low among-species and phylogenetic variation, rather than indicating an effect of adaptation to different environments. Phylogenetic overlap is less pronounced when comparing terrestrial and arboreal taxa; nevertheless, the small sample size imposes substantial limitations.

**Figure 4.**
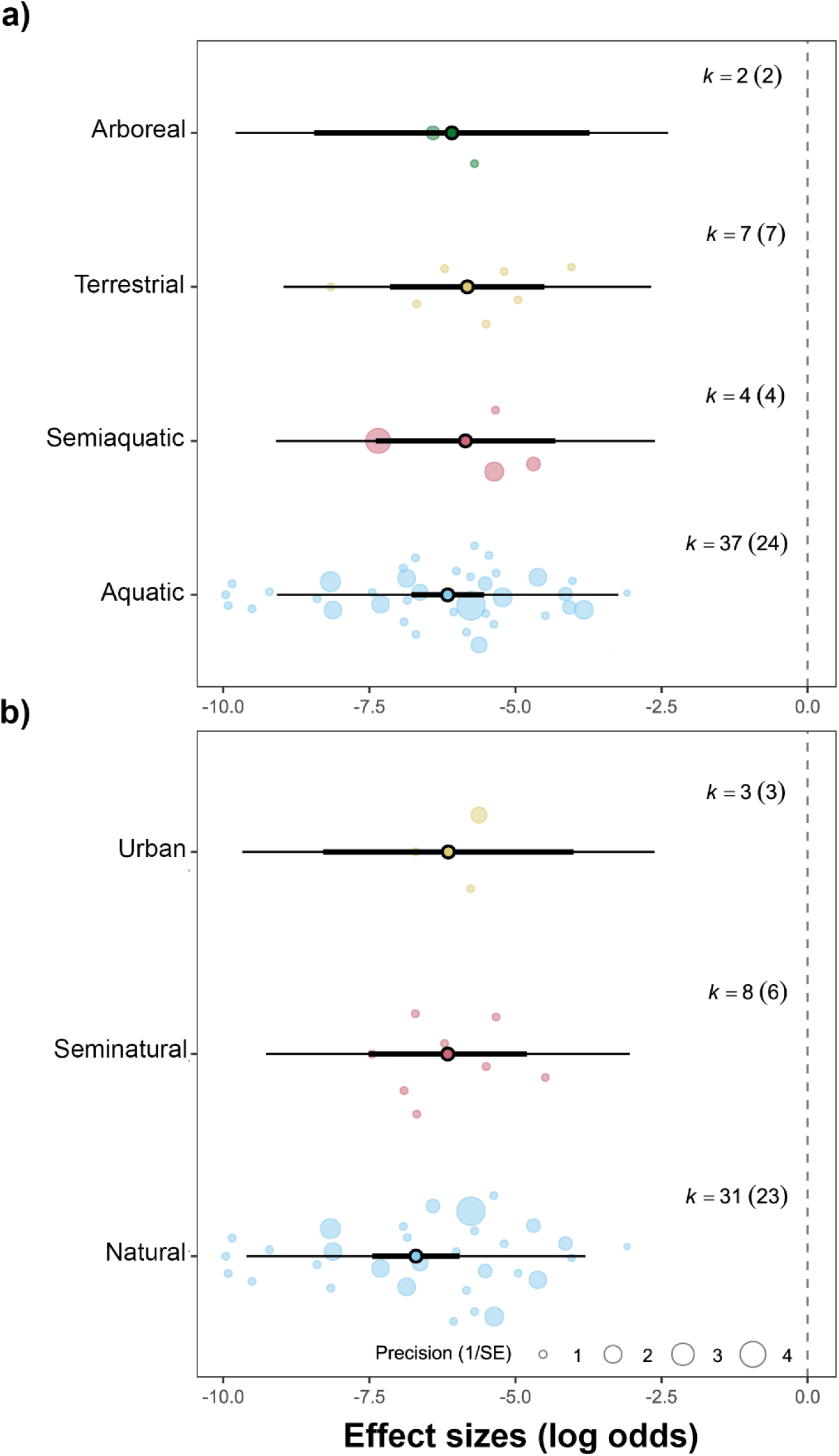
Mean effect sizes by (a) habitat use strategies, and (b) habitat types. Thick horizontal lines represent 95% confidence intervals; thin horizontal lines represent 95% prediction intervals. The point at the centre of each thick line indicates the average effect size. k denotes the number of effect sizes used to calculate the statistics, with the number of studies given in parentheses.

**Figure 5.**
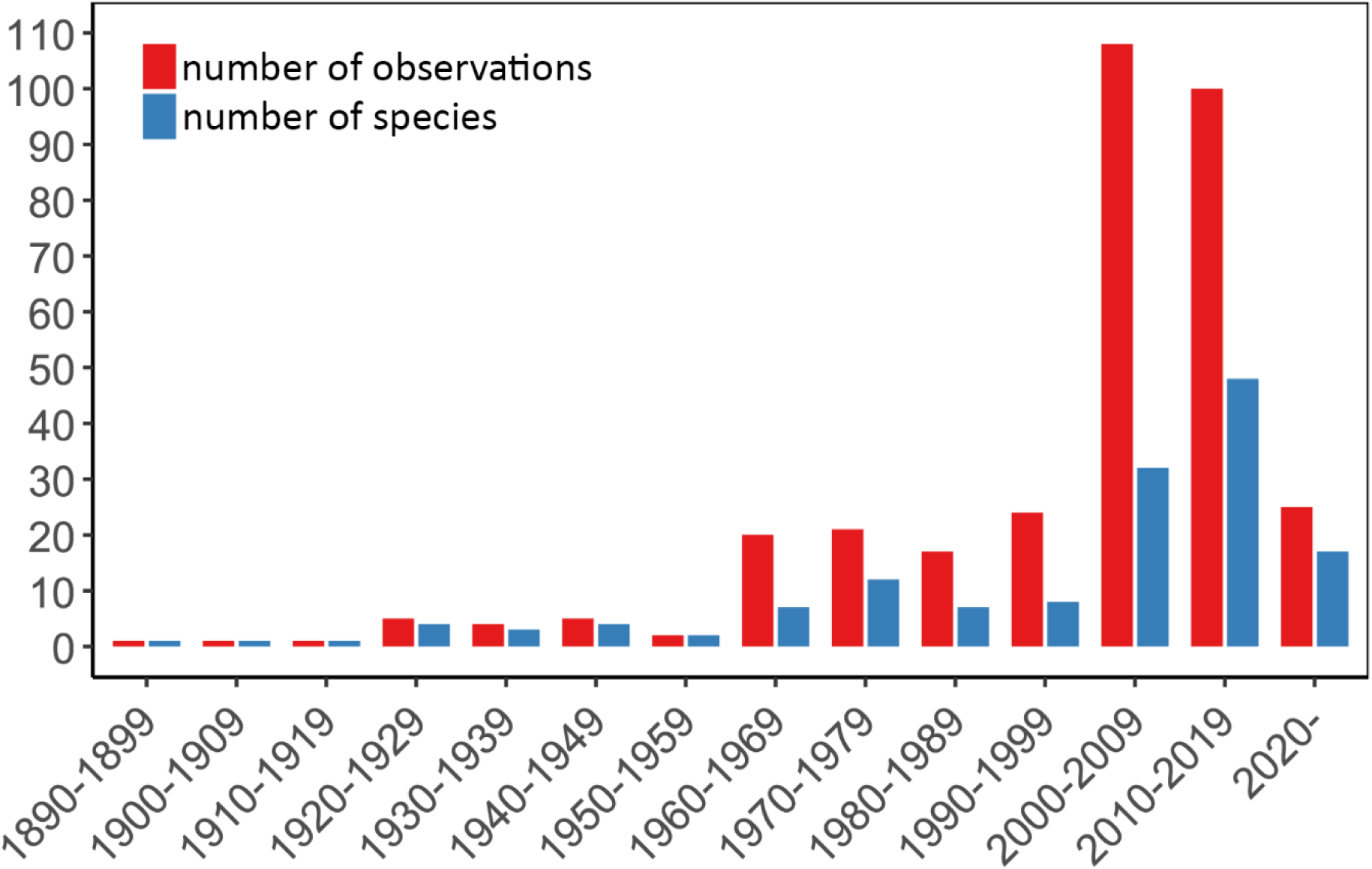
Number of vertebral malformation observations in wild reptiles and the number of affected species according to peer-reviewed and ‘grey’ literature publication records according to the year of observation.

Another aspect to consider is the distribution of various malformation types across habitat use strategies in the complete dataset. Kyphosis was significantly more frequent in aquatic and semiaquatic taxa (79% and 78.8% of records, respectively) than in terrestrial and arboreal taxa (28.6% and 4% of records, respectively). Conversely, scoliosis was more frequent in terrestrial and arboreal taxa (26.8% and 56% of records, respectively) than in aquatic and semiaquatic taxa (9.9% and 6.1% of records, respectively). This pattern aligns with the notion that kyphosis predominantly affects (semi)aquatic turtles, whereas scoliosis is more common in predominantly terrestrial squamates. Nevertheless, this difference is most likely due to substantial differences in the development and structure of the turtle and squamate vertebral column (Houssaye et al., 2010; Nagashima et al., 2012). As the database largely lacks records from (semi)aquatic squamates—e.g., water snakes (*Nerodia* and *Natrix*), water monitors (*Varanus* spp.), and semiaquatic agamids such as the Chinese water dragon (*Physignathus cocincinus*) and the Australian water dragon (*Intellagama lesueurii*)—it remains an open question whether the observed differences are due to habitat use strategy (aquatic *vs.* terrestrial) or rather to phylogenetic, structural, and/or developmental constraints. This issue can only be resolved once more data become available from the aforementioned squamate taxa.

### (3) Does environmental stress increase prevalence?

The prevalence of vertebral deformities did not differ significantly among natural, seminatural, and urbanized habitats (Fig. 4b). Urban environments are recognized as multi-stressor landscapes (Grunst & Grunst, 2024), where individuals are exposed to increased chemical pollution, noise and artificial light pollution, infectious diseases, and poor diet quality (Isaksson, 2015; Putman et al., 2025), compared to natural habitats. Since several of these environmental stressors have been linked to developmental malformations, the absence of a habitat effect on prevalence may seem surprising. However, it is important to note several limitations of the dataset that may have contributed to this negative result. First, I must reiterate that habitat characterization was performed solely by me based on a predefined set of criteria, and it is possible that some misclassifications occurred during the process. Most records originated from habitats identified as (semi)natural, with only three records coming from urbanized habitats (representing 5.8% of the prevalence data), which limits the statistical power to detect weak effects. The low number of prevalence records from urban habitats is somewhat unexpected, as urban reptile populations tend to be well monitored (e.g., Davis & Doherty, 2015; How, Cowan & How 2022; Thongsub et al., 2024).

Importantly, from an ecological perspective, stress is not universal—it is linked to the regulatory capacity of an organism (Koolhaas et al., 2011). Therefore, an environment that is stressful for one taxon may not necessarily pose stress for another. Reptile taxa that have successfully colonized urban habitats may be less sensitive to the stressors typically associated with these environments. While the urban–seminatural–natural trichotomy can serve as a useful proxy for environmental stress, in the absence of detailed data, it was not possible to evaluate the actual level of environmental stress at individual sites. As a result, important differences may remain obscured. The lack of information on potential environmental stressors and the general condition of each habitat represents a significant limitation of the dataset. Only three publications explicitly stated that the habitat was contaminated (e.g., Grey et al., 2003) or referenced prior human disturbance likely to have degraded the environment (e.g., Carvalho et al., 2014). It is essential for future studies and reports to provide information on factors that may act as environmental stressors—such as levels of habitat degradation or contamination—ideally using a standardized framework.

### (4) Potential ecological and behavioural effects

#### (a) Survival and growth

It is commonly assumed that the low prevalence of spinal deformities in wild populations results from the low survivability associated with such conditions. However, this likely depends on (i) whether the malformation is congenital, (ii) the severity of the malformation, and (iii) whether it forms part of a multifaceted pathology. Information about ontogenetic stage was provided in 306 cases, more than half of which (164 records, 53.5%) were adults or subadults, suggesting that in a substantial number of affected individuals does not necessarily impair survival. Nevertheless, it is questionable that in how many of these animals’ condition was congenital. This is very complicated to tell, which is mirrored that only in five cases (3%) was possible to suggest clear congenital origin, while two was acquired post-natal. Juveniles reflect a similar picture: out of 22 records in only six was possible to tell the origin, half of which was congenital, while the other half acquired post-natal. Regarding embryos, hatchlings and neonates (120 records), congenital origin is unambiguous. Several studies— such as Drennen (1990), Grey et al. (2003), Bell et al. (2006), Bárcenas-Ibarra & Maldonado-Gasca (2009), Carvalho (2014), Bárcenas-Ibarra et al. (2015), and Elsey & Stelly (2018)— provide detailed accounts of vertebral malformations affecting embryos, neonates and hatchlings, often in large numbers within the litter. In most of these cases, vertebral deformities were associated with other lethal anomalies incompatible with life. Some studies report affected juveniles died at a very young age (e.g., Progscha & Lehmann, 1970; Wallach & Salmon, 2013), while some studies reported no observable negative effects of vertebral malformations on hatchling, neonates, or juveniles (e.g., Progscha & Lehmann, 1970; Moldowan et al., 2015). If we accept that the majority of spinal deformities are congenital, the above findings suggest that malformations may affect a large number of embryos, of which only a few survive. It is also likely that many malformed individual dies at a young age.

Nevertheless, once an individual reaches adult size, vertebral malformations do not necessarily impair survival. In fact, many reports explicitly state that affected juveniles and (sub)adults were in relatively good condition and health (cf. Carpenter, 1958; Smith & Fitzgerald, 1983; Stuart, 1996; Mitchell & Georgel, 2005; Honarvar et al., 2011; Bernhard et al., 2012; Avila, Medina & Morando, 2013; Brazeau et al., 2013; Datta & Hasan, 2020; Alvarez et al., 2021; Valdez-Villavicencio, León-Serrano & Peralta-García, 2024), sometimes despite the severity of malformation (see Elsey et al., 2017).

Regarding long-term survival and growth patterns, data are available exclusively from long-lived turtle taxa. Information provided by Inwater Research Group Inc. includes loggerhead sea turtle (*Caretta caretta*) and green sea turtle (*Chelonia mydas*) specimens that were recaptured multiple times over a decade. Moldowan et al. (2015) reported a kyphotic painted turtle (*Chrysemys picta marginata*) that evidently lived for at least 17 years, with its condition apparently having no effect on its growth or fitness. Mitchell and Johnson (2014) reported a Florida chicken turtle (*Deirochelys reticularia chrysea*) with kyphoscoliosis that was recaptured three years after the initial encounter and exhibited a seemingly normal growth pattern. Conversely, Harding and Bloomer (1979) and Selman and Jones (2012) found that kyphotic wood turtles (*Glyptemys insculpta*) and ringed map turtles (*Graptemys oculifera*) grew more slowly than expected. Based on these findings, we may conclude that—at least in turtles—vertebral deformities do not hinder normal growth or have only mild effects on it.

#### (b) Escape and predation

Besides direct negative effects on viability, vertebral malformations may increase mortality by making affected individuals more susceptible to predation, mainly due to limited escape capabilities. In the dataset, there were 67 records where information on locomotor abilities was provided, but only 13 studies (19.4%) reported any clear locomotor disorders. These cases involved various, usually severe, types of malformations, suggesting that relatively mild vertebral deformities do not necessarily hinder locomotion, regardless of their type. For example, Mitchell and Georgel (2005) reported a juvenile kyphoscoliotic Eastern fence lizard (*Sceloporus undulatus undulatus*) that actively pursued prey in captivity. Hernández et al. (2024) described a Mesquite lizard (*Sceloporus grammicus*) with thoracic kyphosis and pelvic scoliosis that was observed multiple times moving with speed and agility among rocks.

Several malformed turtles were also found to move and swim normally (e.g., Schachner et al., 2017; Valdez-Villavicencio et al., 2024). In comparison, Avellá Machado and Acosta (2023) mentioned a Darwin’s tree iguana (*Liolaemus darwinii*) that ran without difficulty, although its movement appeared more rigid than that of a healthy individual.

Nevertheless, even if vertebral deformities do not substantially affect locomotor performance, they may compromise the effectiveness of certain antipredator behaviours. For instance, Elsey et al. (2017) reported a severely malformed red-eared slider (*Trachemys scripta elegans*) that appeared healthy and mobile, but its condition prevented complete forelimb retraction and caused abnormal head retraction. Similarly, Hirst et al. (2023) reported a kyphotic spiny chuckwalla (*Sauromalus hispidus*) that seemed to have a limited ability to effectively hide in a burrow. Such limitations are mostly associated with kyphosis and kyphoscoliosis and may particularly affect taxa that rely on crevices and burrows for shelter.

A special type of vertebral deformity also warrants mention in this context: caudal scoliosis, where the malformation is restricted to the tail region, is generally considered unlikely to have severe negative effects on any aspect of the specimen’s life. Records of caudal scoliosis are known only from squamates and accounted for 7.5% of all 134 squamate observations in the database. Although none of these reports mentioned clear locomotor disorders or disadvantages, tail morphology has been shown to influence locomotor performance in lizards (cf. McElroy & Bergmann, 2013), though not in snakes (see Jayne & Bennett, 1989).

Therefore, we may expect some differences between individuals with normal and curled tails, but this issue can only be resolved through targeted measurements.

#### (c) Reproductive outcome

Spinal deformities—especially severe ones—are expected to affect reproductive outcome on multiple levels. First, such malformations may hinder mating success, primarily by impairing courtship and mating behaviours or by making copulation physically impossible. In addition, fertility measures, as well as the number and viability of offspring, may be reduced in malformed mothers. To date, only sporadic explicit or indirect information on the potential effects of spinal deformities on reproductive outcome can be found in the literature.

Importantly, the database indicates that both sexes are affected by spinal deformities, although female records are nearly twice as numerous as male records (124 vs. 76; 62% vs. 38%).

However, since sex was not reported in 151 cases (43% of all records), these proportions may not accurately reflect the true sex ratio.

Although it does not count as mating behaviour *per se*, Norval et al. (2010) mention an adult male scoliotic japalure (*Diploderma chapaense*) that was engaged in a territorial dispute when captured. This observation is important because it represents the only published case providing information on an individual with vertebral deformity expressing specific social behaviour in the wild. Despite not knowing the outcome of this agonistic encounter, it seems not too far-fetched to say that behaviours linked to mating and courtship are expected to be affected alongside locomotor performance—especially in lizards, where active chasing and grabbing of females by males is very common (e.g., Pandav, Shanbhag & Saidapur, 2007; Kim, Kim & Park, 2012). As we have seen, cases with clear indications of locomotor dysfunction, although substantial in number, represent only a minority of records; thus, we may rightly deduce that mating behaviour is relatively rarely affected as well. Nevertheless, beyond actively pursuing their mate, both males and females of several species—especially iguanids—are known to exhibit quite elaborate displays during mating (e.g. Carpenter, 1967; Blanc & Carpenter, 1969), including swinging or curling the tail, behaviours that are surely affected even by mild malformations such as caudal scoliosis. No first-hand information regarding this is available, however.

Regarding copulation success of affected males, we possess no first-hand knowledge. While severe malformations may hinder males across various taxa, in turtles—where kyphosis is seemingly more common—females might be more likely affected, even by relatively mild kyphotic malformations. There is a sole observation that indicates otherwise, or at least suggests that vertebral malformations do not necessarily pose a problem in turtles when it comes to copulation. Burke (1994) reports an adult female Eastern spiny softshell turtle (*Apalone spinifera*) with extreme kyphosis that was gravid and laid a clutch of eggs a few weeks after its capture (oviposition was induced by oxytocin injection). Sadly, the eggs were laid in water, so it was not possible to determine the exact number or fertility of the eggs.

Owens & Knapp (2007) provide valuable information on the number and viability of offspring. They describe a kyphoscoliotic female Andros Island iguana (*Cyclura cyclura cyclura*) that nested in a termite mound. Despite aggressively defending its nest, the nest was excavated, and the eggs inspected. The clutch size of seven eggs was within the normal range for the species; however, the mean egg size was slightly smaller than normal. Despite this, all eggs hatched successfully, and the hatchlings were viable and non-malformed. Owens & Knapp suggest that the spinal curvatures may have resulted in a restricted cloacal opening and constrained egg width, while a reduced abdominal cavity may have caused the below-average egg mass. The existence of such additional effects and their distribution across various taxa could be relatively easily confirmed using modern imaging techniques on malformed females.

## V. FUTURE DIRECTIONS

Reports of spinal deformities in wild reptiles were sporadic throughout the last century. However, over the past 25 years, notes and short communications documenting individuals with such malformations have become increasingly common, with nearly ten publications per year on average. This rise has enabled the production of some thematic syntheses (e.g., Rhodin, Pritchard & Mittermeier, 1984; Bateman et al., 2022), although these studies offer limited insight into ecological implications. Most publications remain restricted to single-case observations that include morphological measurements and, occasionally, data on occurrence frequency. Another issue is that, despite the growing number of peer-reviewed papers, spinal deformities are still often regarded as mere curiosities—even by professionals. As a result, a substantial number of observations likely remain unpublished or are available only in grey literature, where they tend to go unnoticed.

Here, I compiled a global database of recorded spinal deformities in wild reptiles. Complemented by a phylogenetic meta-analysis and informed by the limited known effects of kyphosis and scoliosis on ecology and life history, this work is intended to spark a lively discussion on the potential ecological implications of spinal deformities. In addition, the present work aims to provide a blueprint for simple approaches to future observations, in order to enhance their ecological relevance and our overall understanding of the ecological effects of spinal deformities in wild reptiles.

i. Observations of spinal deformities—even single cases—are not mere curiosities; they provide important scientific information. Therefore, it is essential to publish them on scientific platforms, preferably in peer-reviewed herpetological journals. In many cases, malformed animals—especially turtles—are observed from a distance and cannot be captured; in such instances, photographic documentation of the specimen(s) is the minimum expected standard.
ii. Information on the frequency of occurrences within populations and habitat characteristics should be provided. These data can help determine how common spinal deformities are in a given population and identify potential environmental factors—particularly environmental pollution—that may influence their occurrence. In the absence of first-hand information on abundance and population density (which are needed to calculate prevalence), future studies should consider alternative approaches to obtain these data, such as contacting conservation agencies or national parks. Alternatively, online community science data repositories—such as iNaturalist—may provide information on local population abundance. Regarding environmental pollution and habitat degradation, as much information as possible should be collected to identify the most significant environmental stressors. However, if no specific data are available, the level of urbanization may serve as a useful—albeit limited—proxy.
iii. Provide information on morphology (snout-vent-length and tail length for squamates, carapace length, carapace width and carapace height for turtles), body mass, ontogenetic stage and sex.
iv. The exact type of malformation should be confirmed, along with a detailed description of the affected regions of the spine, the number of curvatures, and any additional abnormalities. Since visual observation is not always sufficient, the use of modern imaging techniques— such as radiography and CT scanning—is strongly encouraged. These methods are valuable for detecting concurrent physical abnormalities that may influence life-history traits.
v. The collection of blood samples is encouraged. Blood cultures offer a relatively simple and minimally invasive method for diagnosing active bacterial osteomyelitis and osteoarthritis. This approach may help advance our understanding of the aetiology of vertebral malformations in wild reptiles.
vi. Testing potential behavioural, locomotor, and foraging efficiency limitations associated with spinal deformities is essential for assessing the extent to which these malformations are detrimental to the individual. Such investigations can clarify their role in shaping behavioural ecology and provide insight into how spinal deformities affect the survival of various reptile taxa.

## VI. CONCLUSIONS

1. This review presents the most comprehensive documentation and discussion of spinal deformities—primarily kyphosis, scoliosis, and kyphoscoliosis—in wild reptiles to date. Spinal deformities have been recorded in reptiles for over 100 years, occurring in 103 species across 24 of the currently recognized 94 reptile families. With the exception of tuataras and amphisbaenians, all major reptile clades are affected, with particularly high numbers of cases reported in Testudines.
2. It is safe to say that published reports significantly underrepresent the true number of spinal deformity cases in wild reptiles. Observation frequency appears to be more strongly influenced by research focus and taxonomic popularity than by actual incidence, while the geographical distribution of reports likely reflects the presence of professional herpetologists and amateur enthusiasts rather than underlying species richness. As shown previously, differences in language, culture, and scientific practices may hinder both the publication and accessibility of certain records. Therefore, reporting observations of spinal deformities in reptiles—preferably in peer-reviewed herpetological journals—is highly encouraged.
3. Meta-analytic results revealed no significant effect of phylogeny, habitat use strategy, or habitat type. However, the absence of detectable effects is likely due to limitations in the available data. Notably, the significant overrepresentation of (semi)aquatic turtles may contribute to low phylogenetic variation. Additional data from squamate taxa—particularly those with (semi)aquatic lifestyles—are needed to better assess potential phylogenetic differences. Future research should also place greater emphasis on characterizing the frequency of occurrences within populations and on providing detailed habitat descriptions.
4. Spinal deformations have the potential to severely impact individual ecology and life history traits, particularly escape and survival, growth, and reproductive output. However, direct data are scarce, and current inferences can only be drawn from isolated, specific cases.
5. To enhance ecological relevance, future studies should adopt a more detailed and preferably standardized reporting approach that includes information on occurrence frequency, habitat characteristics, morphometry, the use of imaging techniques, and assessments of potential limitations in behaviour, locomotion, and foraging efficiency.

## VII: ACKNOWLEDGEMENTS

I am grateful to many individuals who provided personal observation data and/or photographs: Rita Babos, Henrik Bringsøe, Dennis Caldwell, Petrișor Mădălina, Doru Panatinescu, Inwater Research Group Inc., Ivo Peranic, Octavio Jiménez Robles, Stacey Schenkel, Jackson D. Shedd, Yoel E. Stuart, Bálint Üveges and Mark Witwer. My sincere thanks also go to Wolfgang Böhme, Mike Dloogatch, Nicolas Dowdy, John Iverson, Renata Platenberg, Anders G.J. Rhodin, Réka Sebestyén, James N. Stuart, Judit Vörös and Susan Walls for providing essential bibliography for this project. I appreciate Gábor Herczeg for his impactful comments during preliminary analysis of the dataset. This project has been funded by the HUN-REN Hungarian Research Network and supported by the János Bolyai Research Scholarship of the Hungarian Academy of Sciences.

## XI. SUPPORTING INFORMATION

**Fig. S1.**
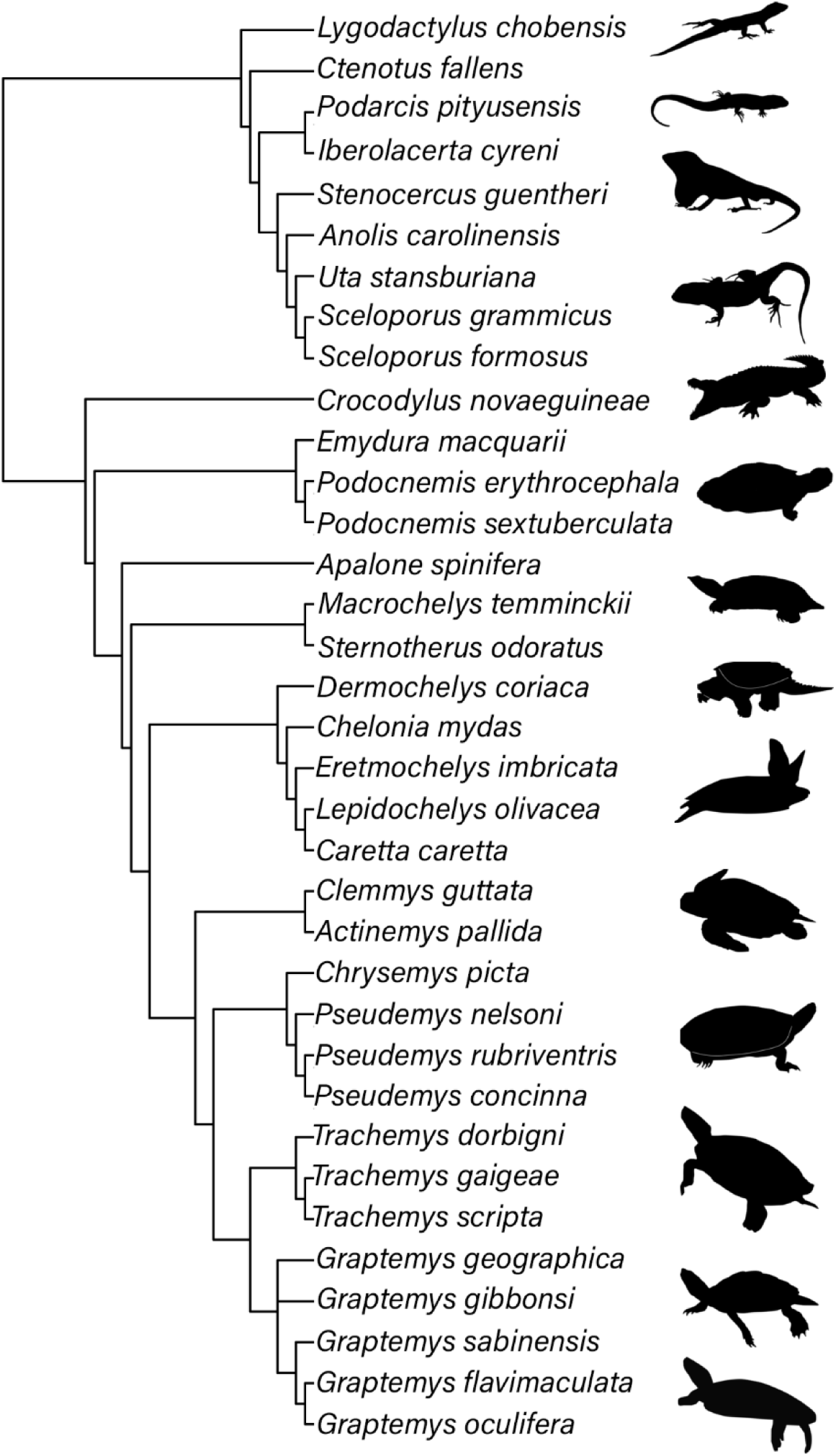
Graphical summary of phylogenetic distribution of 35 species included in the meta- analyses. Note that, for compatibility with the Open Tree of Life database, the Suwannee snapping turtle (*Macrochelys suwanniensis*) was treated as a synonym of the alligator snapping turtle (*M. temminckii*). The tree was created using the ‘rotl’ package (Michonneau, Brown, & Winter, 2016) in R. Silhouettes depicting selected taxa obtained from phylopic.org.

**Fig. S2.**
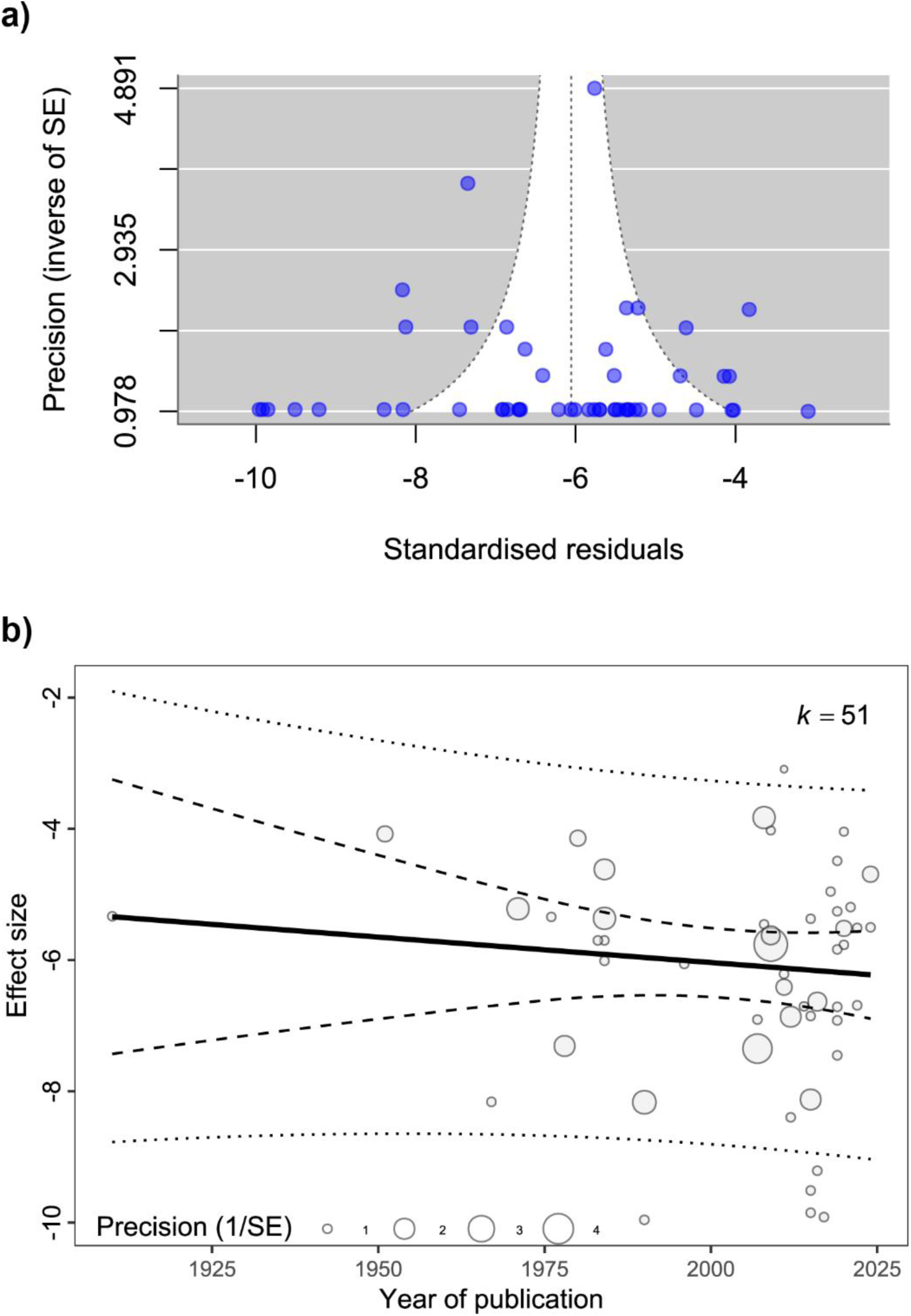
Funnel plot and relationships between effect sizes and publication year. a) Funnel plot using effect size and its inverse standard error; b) the relationship between effect sizes and publication year. In b) circle sizes are scaled accordingly to precision, and k represents the number of effect sizes. Fitted regression line is shown as a straight line, and 95% confidence and prediction intervals are shown as dashed and dotted lines, respectively.

**Supplementary Table S1.** Occurrence and prevalence of spinal deformations in the currently known families of reptiles. Families are consistent with the most recent classifications from The Reptile Database (Uetz et al., 2025; http://www.reptile-database.org, accessed May 24th 2025). Numbers in parentheses are the number of species observed from the literature search to have spinal deformities and the number of known species in the family.

| Order | Family | Species | Abnormality | Prevalence (%) | Reference |
| --- | --- | --- | --- | --- | --- |
| Crocodylia | Crocodylidae (2/17) | *Crocodylus acutus* | scoliosis |  | Rossmann, 2001 |
|  |  | *Crocodylus novaeguineae* | kyphosis | 0.47 | Montague 1984 |
|  | Gavialidae (0/2) | - | - | - | - |
|  | Alligatoridae (1/8) | *Alligator mississippiensis* | kyphoscoliosis | - | Elsey & Stelly, 2018 |
| Rhynchocephalia | Sphenodontidae (0/1) | - | - | - | - |
| Squamata | Agamidae (7/602) | *Agama anchietae* | scoliosis | - | Grogan, 1976 |
|  |  | *Agama bibronii* | scoliosis | - | Ahboucha & Gamrani, 2001 |
|  |  | *Calotes goetzi* | kyphoscoliosis | - | Henrik Bringsøe; personal communication |
|  |  | *Calotes versicolor* | scoliosis | - | Henrik Bringsøe; personal communication |
|  |  | *Calotes versicolor* | kyphosis | - | Datta & Hasan, 2020 |
|  |  | *Diploderma chapaense* | scoliosis | - | Norval et al., 2010 |
|  |  | *Trapelus mutabilis* | kyphoscoliosis | - | Böhme, 1970 |
|  | Chamaeleonidae (1/234) | *Furcifer pardalis* | kyphoscoliosis | - | Gehring, 2009 |
|  | Corytophanidae (0/11) | - | - | - | - |
|  | Crotaphytidae (0/12) | - | - | - | - |
|  | Anolidae (4/435) | *Anolis carolinensis* | scoliosis | 0.16 | Stuart, 2011; online, personal communication |
|  |  | *Anolis cybotes* | kyphoscoliosis | - | Ramírez Tiburcio et al., 2021 |
|  |  | *Anolis sagrei* | scoliosis | - | Campbell, 1996 |
|  |  | *Anolis sagrei* | scoliosis | - | Losos, 2014; online |
|  |  | *Anolis sericeus* | kyphoscoliosis | - | Ortiz-Medina & Valdez-Villavicencio, 2016 |
|  | Hoplocercidae (0/22) | - | - | - | - |
|  | Iguanidae (2/45) | *Sauromalus hispidus* | kyphosis | - | Hirst et al., 2023 |
|  |  | *Cyclura cyclura* | kyphoscoliosis | - | Owens & Knapp, 2007 |
|  | Leiocephalidae (0/30) | - | - | - | - |
|  | Leiosauridae (0/36) | - | - | - | - |
|  | Liolaemidae (6/342) | *Liolaemus baguali* | kyphosis | - | Feltrin et al., 2009 |
|  |  | *Liolaemus darwini* | kyphoscoliosis | - | Avellá Machado & Acosta, 2023 |
|  |  | *Liolaemus fittkaui* | kyphoscoliosis | - | Octavio Jiménez-Robles; personal communication |
|  |  | *Liolaemus huacahuascianus* | kyphosis | - | Halloy & Laurent, 1988 |
|  |  | *Liolaemus koslowskyi* | kyphoscoliosis | - | Avila et al., 2013 |
|  |  | *Liolaemus petrophilus* | scoliosis | - | Frutos et al., 2006 |
|  | Opluridae (0/8) | - | - | - | - |
|  | Phrynosomatidae (9/170) | *Phrynosoma douglasii* | kyphosis? | - | Jackson Shedd; personal communication |
|  |  | *Sceloporus formosus* | kyphoscoliosis | 1.72 | Castillo-Juárez et al., 2020 |
|  |  | *Sceloporus graciosus* | kyphoscoliosis | - | Valdez-Villavicencio et al., 2016 |
|  |  | *Sceloporus grammicus* | kyphoscoliosis | 0.41 | Hernández et al., 2024 |
|  |  | *Sceloporus magister* | kyphoscoliosis | - | Dennis Caldwell; personal communication |
|  |  | *Sceloporus marmoratus* | kyphoscoliosis | - | Chávez-Cisneros & Lazcano, 2012 |
|  |  | *Sceloporus torquatus* | kyphosis | - | Pérez-Delgadillo et al., 2015 |
|  |  | *Sceloporus undulatus* | kyphoscoliosis | - | Mitchell & Georgel, 2005 |
|  |  | *Uta stansburiana* | scoliosis | - | Stacey Schenkel; personal communication |
|  |  | *Uta stansburiana* | scoliosis | 0.03 | Tinkle, 1967 |
|  | Polychrotidae (0/8) | - | - | - | - |
|  | Tropiduridae (2/149) | *Plica umbra* | kyphoscoliosis | - | Carvalho et al., 2014 |
|  |  | *Stenocercus guentheri* | kyphoscoliosis | 0.69 | Ramírez-Jaramillo, 2018 |
|  | Gekkonidae (3/1680) | *Lygodactylus chobiensis* | scoliosis | 0.33 | Simbotwe, 1983 |
|  |  | *Phelsuma grandis* | rhoecosis | - | Caceres et al., 2024 |
|  |  | *Phelsuma modesta* | kyphoscoliosis | - | Gehring et al. 2010 |
|  | Carphodactylidae (0/34) | - | - | - | - |
|  | Diplodactylidae (0/202) | - | - | - | - |
|  | Eublepharidae (0/48) | - | - | - | - |
|  | Phyllodactylidae (0/168) | - | - | - | - |
|  | Sphaerodactylidae (0/238) | - | - | - | - |
|  | Pygopodidae (0/47) | - | - | - | - |
|  | Cordylidae (0/69) | - | - | - | - |
|  | Gerrhosauridae (0/39) | - | - | - | - |
|  | Scincidae (4/1785) | *Ctenotus fallens* | scoliosis | 0.12 | Bateman et al., 2022 |
|  |  | *Lampropholis guichenoti* | scoliosis | - | Philip Bateman; personal communication |
|  |  | *Marisora brachypoda* | kyphosis | - | Arrivillaga & Brown, 2019 |
|  |  | *Trachylepis seychellensis* | kyphosis | - | Petrișor Mădălina; personal communication |
|  | Xantusiidae (0/38) | - | - | - | - |
|  | Alopoglossidae (0/32) | - | - | - | - |
|  | Gymnophthalmidae<br>(0/297) | - | - | - | - |
|  | Lacertidae (5/387) | *Iberolacerta cyreni* | kyphosis | 0.55 | Horváth et al., 2021 |
|  |  | *Lacerta agilis* | kyphoscoliosis | - | Geuss, 1966 |
|  |  | *Lacerta agilis* | scoliosis | - | Gordeev et al., 2021 |
|  |  | *Podarcis pityusensis* | kyphosis | 0.2 | Garin-Barrio et al., 2011 |
|  |  | *Psammodromus*<br>*hispanicus* | scoliosis? | - | Doru Panaitescu; personal<br>communication |
|  |  | *Timon Lepidus* | kyphosis | - | Berdún & Bisbal-Chinesta, 2018 |
|  |  | *Timon lepidus* | kyphosis | - | Martínez-Rodríguez, 2017; online |
|  | Teiidae (1/173) | *Aspidoscelis tigris* | scoliosis | - | Heyborne, 2021 |
|  | Anguidae (0/88) | - | - | - | - |
|  | Diploglossidae (0/90) | - | - | - | - |
|  | Xenosauridae (0/14) | - | - | - | - |
|  | Amphisbaenidae (0/184) | - | - | - | - |
|  | Bipedidae (0/3) | - | - | - | - |
|  | Blanidae (0/7) | - | - | - | - |
|  | Cadeidae (0/2) | - | - | - | - |
|  | Rhineuridae (0/1) | - | - | - | - |
|  | Trogonophidae (0/6) | - | - | - | - |
|  | Helodermatidae (0/5) | - | - | - | - |
|  | Lanthanotidae (0/1) | - | - | - | - |
|  | Varanidae (0/88) | - | - | - | - |
|  | Shinisauridae (0/1) | - | - | - | - |
|  | Dibamidae (0/28) | - | - | - | - |
|  | Acrochordidae (0/3) | - | - | - | - |
|  | Anomochilidae (0/3) | - | - | - | - |
|  | Cylindrophiiidae (0/15) | - | - | - | - |
|  | Uropeltidae (0/73) | - | - | - | - |
|  | Loxocemidae (0/1) | - | - | - | - |
|  | Pythonidae (0/38) | - | - | - | - |
|  | Xenopeltidae (0/3) | - | - | - | - |
|  | Boidae (1/67) | *Sanzinia madagascariensis* | kyphoscoliosis<br>, scoliosis | - | Progscha & Lehmann, 1970 |
|  | Colubridae (7/2156) | *Arizona elegans* | kyphoscoliosis | - | Alvarez et al., 2021 |
|  |  | *Coronella austriaca* | lordosis | - | Najbar et al., 2022 |
|  |  | *Nerodia sipedon* | scoliosis? | - | Wallach & Salmon, 2013 |
|  |  | *Oxybelis microphthalmus* | scoliosis | - | Nieto-Toscano & Martínez-Coronel, 2023 |
|  |  | *Pantherophis gloydi* | kyphosis | - | Brazeau et al., 2013 |
|  |  | *Thamnopsis radix* | rhoecosis | - | Smith & Fitzgerald, 1983 |
|  |  | *Thamnopsis sirtalis* | scoliosis | - | Gray et al., 2003 |
|  | Atractaspididae (0/68) | - | - | - | - |
|  | Cyclocoridae (0/8) | - | - | - | - |
|  | Micrelapidae (0/6) | - | - | - | - |
|  | Lamprophiidae (0/93) | - | - | - | - |
|  | Prosymnidae (0/19) | - | - | - | - |
|  | Psammodynastidae (0/2) | - | - | - | - |
|  | Psammophiidae (0/57) | - | - | - | - |
|  | Pseudaspididae (0/2) | - | - | - | - |
|  | Pseudoxyrhophiidae<br>(0/89) | - | - | - | - |
|  | Elapidae (0/416) | - | - | - | - |
|  | Anomalepididae (0/23) | - | - | - | - |
|  | Gerrhopilidae (0/29) | - | - | - | - |
|  | Typhlopidae (0/282) | - | - | - | - |
|  | Leptotyphlopidae (0/146) | - | - | - | - |
|  | Xenotyphlopidae (0/1) | - | - | - | - |
|  | Aniliidae (0/1) | - | - | - | - |
|  | Bolyeriidae (0/2) | - | - | - | - |
|  | Homalopsidae (0/60) | - | - | - | - |
|  | Pareidae (0/45) | - | - | - | - |
|  | Tropidophiidae (0/37) | - | - | - | - |
|  | Viperidae (5/405) | *Bothrops jararaca* | kypholordosis,<br>kyphoscoliosis<br>, kyphosis,<br>lordoscoliosis,<br>lordosis,<br>rhoecosis,<br>scoliosis | - | Carvalho, 2014 |
|  |  | *Crotalus durissus* | kyphoscoliosis<br>, kyphosis,<br>lordosis,<br>rhoecosis,<br>scoliosis | - | Carvalho, 2014 |
|  |  | *Vipera ammodytes* | scoliosis | - | Ivo Peranic; personal<br>communication |
|  |  | *Vipera graeca* | scoliosis? | - | Bálint Üveges; personal<br>communication |
|  |  | *Vipera ursinii moldavica* | scoliosis? |  | Bálint Üveges; personal communication |
|  |  | *Vipera ursinii rakosiensis* | scoliosis?<br>kyphosis? |  | Bálint Üveges; personal communication |
|  | Xenodermidae (0/37) | - | - | - | - |
|  | Xenophidiidae (0/2) | - | - | - | - |
| Testudines | Emydidae (27/58) | *Actinemys pallida* | kyphosis | 0.91 | Valdez-Villavicencio et al., 2024 |
|  |  | *Chrysemys picta* | kyphosis | 0.54 | Ernst, 1971 |
|  |  | *Chrysemys picta* | kyphosis | - | Pavalko, 1986 |
|  |  | *Chrysemys picta* | kyphosis | - | Thomas Spence; online |
|  |  | *Chrysemys picta bellii* | scoliosis | 1.56 | MacCulloch, 1980 |
|  |  | *Chrysemys picta bellii* | kyphosis | 0.23 | Stuart, 1996 |
|  |  | *Chrysemys picta bellii* ×<br>*marginata* | kyphosis | - | Necker, 1940 |
|  |  | *Chrysemys picta*<br>*marginata* | kyphosis | - | Ernst, 1971 |
|  |  | *Chrysemys picta marginata* | kyphosis | 0.11 | Moldowan et al., 2015 |
|  |  | *Chrysemys picta marginata* | kyphosis | - | Werner, 1959 |
|  |  | *Chrysemys picta picta* | lordosis | 0.12 | Mitchell, 2014 |
|  |  | *Trachemys yaquia* | kyphosis | - | Plymale et al., 1978 |
|  |  | *Clemmys guttata* | kyphosis | 0.48 | Ernst, 1976 |
|  |  | *Glyptemys insculpta* | kyphosis | - | Harding and Bloomer, 1979 |
|  |  | *Glyptemys insculpta* | scoliosis | - | Jonathan Mays; online |
|  |  | *Deirochelys reticularia chrysea* | kyphoscoliosis | - | Mitchell & Johnson, 2014 |
|  |  | *Emys orbicularis* | kyphosis? | - | Rita Babos; online, personal communication |
|  |  | *Emys orbicularis* | lordosis | - | Valdeón et al., 2020 |
|  |  | *Graptemys flavimaculata* | kyphosis | 0.09 | Mitchell et al., 2019 |
|  |  | *Graptemys flavimaculata* | scoliosis | - | Selman, 2017 |
|  |  | *Graptemys geographica* | kyphoscoliosis | - | Bennett & Litzgus, 2014 |
|  |  | *Graptemys geographica* | kyphosis | 1.11 | Mitchell et al., 2019 |
|  |  | *Graptemys geographica* | kyphosis | 0.06 | Mitchell et al., 2019 |
|  |  | *Graptemys geographica* | kyphosis | - | Mitchell et al., 2019 |
|  |  | *Graptemys geographica* | kyphosis | 0.12 | Mitchell et al., 2019 |
|  |  | *Graptemys gibbonsi* | kyphosis | 0.29 | Mitchell et al., 2019 |
|  |  | *Graptemys oculifera* | kyphosis | 0.1 | Selman & Jones, 2012 |
|  |  | *Graptemys ouachitensis* | kyphosis | - | Carpenter, 1958 |
|  |  | *Graptemys sabinensis* | kyphosis | 0.46 | Loque Jr. et al., 2015 |
|  |  | *Graptemys sabinensis* | kyphoscoliosis | 0.46 | *Graptemys sabinensis* |
|  |  | *Graptemys versa* | kyphosis | - | Franklin, 2021 |
|  |  | *Malaclemys terrapin*<br>*centrata* | lordosis?,<br>scoliosis? | - | Hildebrand, 1930 |
|  |  | *Malaclemys terrapin*<br>*centrata* | kyphosis,<br>lordosis | - | Selman, 2019 |
|  |  | *Podocnemis erythrocephala* | kyphosis | 0.02 | Bernhard et al., 2012 |
|  |  | *Pseudemys concinna suwanniensis* | kyphoscoliosis | 0.13 | Mitchell & Johnston, 2016 |
|  |  | *Pseudemys gorzugi* | kyphosis | - | Waldon et al., 2018 |
|  |  | *Pseudemys gorzugi* | kyphoscoliosis<br>? | - | Zymonas, 2009 |
|  |  | *Pseudemys nelsoni* | kyphoscoliosis | 0.31 | Jackson & Zappalorti, 2020 |
|  |  | *Pseudemys peninsularis* | kyphosis | - | Donini et al., 2020 |
|  |  | *Pseudemys peninsularis* | kyphosis | - | Mark Witwer; online |
|  |  | *Pseudemys rubriventris* | kyphosis | 0.4 | Jackson & Zappalorti, 2020 |
|  |  | *Terrapene carolina* | scoliosis | - | Palis, 2017 |
|  |  | *Terrapene ornata* | kyphoscoliosis | - | Fox, 1941 |
|  |  | *Terrapene triunguis* | kyphosis | - | Black, 1976 |
|  |  | *Trachemys dorbigni* | kyphosis | 1.75 | Bujes, 2009 |
|  |  | *Trachemys gaigeae*<br>*gaigeae* | kyphoscoliosis<br>? | 0.43 | Stuart & Painter, 2008 |
|  |  | *Trachemys gaigeae*<br>*gaigeae* | kyphosis | 2.13 | Stuart & Painter |
|  |  | *Trachemys scripta elegans* | kyphoscoliosis | - | Elsey et al., 2017 |
|  |  | *Trachemys scripta elegans* | kyphosis | 0.06 | Tucker et al., 2007 |
|  |  | *Trachemys scripta elegans* | kyphosis | - | Carr, 1952 |
|  |  | *Trachemys scripta elegans* | kyphoscoliosis | - | Enge et al., 2022 |
|  | Testudinidae (0/47) |  |  |  |  |
|  | Geoemydidae (0/75) |  |  |  |  |
|  | Platysternidae (0/1) |  |  |  |  |
|  | Carettochelydae (0/1) |  |  |  |  |
|  | Trionychidae (7/36) | *Apalone ferox* | kyphosis | - | Nixon & Smith, 1949 |
|  |  | *Apalone ferox* | kyphosis | - | Pritchard, 2008 |
|  |  | *Apalone ferox* | kyphosis | - | Taylor & Mendyk, 2017 |
|  |  | *Apalone mutica* | kyphosis | - | Smith, 1947 |
|  |  | *Apalone mutica mutica* | kyphosis | - | Webb, 1962 |
|  |  | *Apalone spinifera* | kyphosis | - | Burke, 1994 |
|  |  | *Apalone spinifera* | kyphosis | - | Cahn, 1937 |
|  |  | *Apalone spinifera* | kyphosis | - | Smith, 1947 |
|  |  | *Apalone spinifera* | kyphosis | - | Webb, 1962 |
|  |  | *Apalone spinifera emoryi* | kyphosis | - | Smith, 1947 |
|  |  | *Apalone spinifera emoryi* | kyphosis | - | Stuart, 1996 |
|  |  | *Apalone spinifera emoryi* | kyphosis | - | Webb, 1962 |
|  |  | *Apalone spinifera hartwegi* | kyphosis | - | Webb, 1962 |
|  |  | *Apalone spinifera spinifera* | kyphosis | 1.67 | Neill, 1951 |
|  |  | *Apalone spinifera spinifera* | kyphosis | - | Webb, 1962 |
|  |  | *Apalone spinifera spinifera* | kyphosis | - | White & Murphy, 1972 |
|  |  | *Lissemys punctata punctata* | kyphosis | - | Duda & Gupta, 1977 |
|  |  | *Palea steindachneri* | kyphosis | - | Gressit, 1936c |
|  |  | *Pelodiscus sinensis* | kyphosis | - | Vogt, 1922 |
|  |  | *Trionyx triunguis* | kyphosis | - | Mertens, 1940 |
|  |  | *Trionyx triunguis* | lordosis | - | Pritchard, 2008 |
|  | Chelydridae (3/5) | *Chelydra serpentina* | kyphosis,<br>scoliosis | - | Bell et al., 2006 |
|  |  | *Chelydra serpentina* | kyphosis | - | Chan, 1937 |
|  |  | *Chelydra serpentina* | kyphoscoliosis | - | Schachner et al., 2017 |
|  |  | *Chelydra serpentina* | kyphosis | - | Wilhoft, 1980 |
|  |  | *Macrochelys suwanniensis* | kyphosis | 0.4 | Enge et al., 2022 |
|  |  | *Macrochelys suwanniensis* | kyphosis | - | Jake Scott; online |
|  |  | *Macrochelys temminckii* | kyphosis | 0.52 | Pearson et al., 2019 |
|  |  | *Macrochelys temminckii* | kyphosis | - | Pritchard, 1989 |
|  | Dermatemydidae (0/1) |  |  |  |  |
|  | Kinosternidae (1/33) | *Sternotherus odoratus* | kyphosis | 0.1 | Iverson, 2007 |
|  |  | *Sternotherus odoratus* | kyphosis | - | Nixon & Smith, 1949 |
|  |  | *Sternotherus odoratus* | kyphosis | - | Saumure, 2001 |
|  | Cheloniidae (4/6) | *Caretta caretta* | scoliosis | 0.48 | Coker, 1910 |
|  |  | *Caretta caretta* | kyphosis | 0.005 | Drennen, 1990 |
|  |  | *Caretta caretta* | scoliosis | 0.03 | Drennen, 1990 |
|  |  | *Caretta caretta* | kyphosis,<br>kyphosis?,<br>lordosis | - | Inwater Research Group Inc.;<br>personal communication |
|  |  | *Caretta caretta* | scoliosis | - | Nardini et al., 2016 |
|  |  | *Caretta caretta* | kyphosis | - | Orós et al., 2005 |
|  |  | *Caretta caretta* | kyphoscoliosis<br>? | - | Wilson et al., 2022 |
|  |  | *Chelonia mydas* | kyphosis | - | Inwater Research Group Inc.;<br>personal communication |
|  |  | *Chelonia mydas* | kyphosis | - | Moosnipol Educación Ambiental y Conservación; online |
|  |  | *Chelonia mydas* | kyphosis | 0.98 | Rhodin et al., 1984 |
|  |  | *Chelonia mydas* | lordosis | 0.25 | Rhodin et al., 1984 |
|  |  | *Eretmochelys imbricata* | kyphosis | 0.005 | Bárcanes-Ibarra et al., 2015 |
|  |  | *Eretmochelys imbricata* | kyphosis | - | Inwater Research Group Inc.; personal communication |
|  |  | *Lepidochelys olivacea* | kyphosis | 0.31 | Bárcanes-Ibarra & Maldonado-Gasca, 2009 |
|  |  | *Lepidochelys olivacea* | kyphosis | 0.007 | Bárcanes-Ibarra et al., 2015 |
|  |  | *Lepidochelys olivacea* | scoliosis | 0.03 | Bárcanes-Ibarra et al., 2015 |
|  |  | *Lepidochelys olivacea* | scoliosis | 0.005 | Bárcanes-Ibarra et al., 2015 |
|  |  | *Lepidochelys olivacea* | kyphosis | 0.33 | Rhodin et al., 1984 |
|  | Dermochelyidae (1/1) | *Dermochelys coriacea* | kyphosis | 0.07 | Fretey, 1978 |
|  |  | *Dermochelys coriacea* | kyphosis | 4.34 | Honarvar et al., 2011 |
|  | Chelidae (2/67) | *Elseya irwini* | kyphoscoliosis | - | Turner, 2018 |
|  |  | *Emydura macquarii*<br>*krefftii* | kyphosis | 0.36 | Trembath, 2009 |
|  | Pelomedusidae (0/27) | - | - | - | - |
|  | Podocnemididae (1/8) | *Podocnemis*<br>*sextuberculata* | kyphosis | 0.01 | Perrone et al., 2016 |

## X. DATA AVAILABILITY

Data and detailed code for the analyses can be found at the Open Science Framework (OSF): https://osf.io/agnzt/?view_only=6c5845bde2f54496af577165771ccb6b

